# BirdCODE: Detecting bird communication at scale

**DOI:** 10.64898/2026.07.31.742086

**Authors:** Anthony Fine, Benjamin Hoffman, David Robinson, Marius Miron, Milad Alizadeh, Emmanuel Chemla, Maddie Cusimano, Logan S. James, Sara Keen, Emma Matthies, Gagan Narula, Inês Nolasco, Matthieu Geist

## Abstract

Deep learning-based animal sound identification is regularly applied to large audio datasets for ecological monitoring and citizen science, but existing methods lack the fine temporal resolution required to extract insights into animal communication from these same datasets. Here we introduce Bird Communication Detector (BirdCODE), a deep learning model that detects and classifies the vocalizations of over 9000 bird species with precise temporal boundaries, a several hundredfold increase the number of species over previous bioacoustic sound event detection models. In extensive benchmarking, BirdCODE achieves state-of-the-art performance in detection and classification of bird sounds. Applying BirdCODE to 1.3M citizen-science recordings, we present four case studies of how BirdCODE-computed sound event boundaries can be used to carry out phylogenetic analyses, to describe geographic and temporal variation in acoustic communication, and to characterize cross-species interactions. Together, these demonstrate how BirdCODE can enable large-scale, data-driven studies of bird communication. Model code, weights, and predictions are publicly available.

---

Passive acoustic monitoring and citizen science platforms have produced millions of audio recordings of wild animals [43, 12, 31]. The vocalizations captured in these recordings encode information about species occurrence at particular times and locations, thereby enabling largescale ecological monitoring programs [16]. These recordings also capture rich details of animal communication beyond species identity — how signals are used and how they vary with geography, time, habitat, and external stimuli — opening the door to studying the evolutionary and behavioral diversity of communication at a global scale.

However, to pursue this opportunity, it is necessary to first quantify the communication behavior within the recordings at a fine temporal scale, at a minimum by labeling the species, onset time, and offset time of each vocalization (together, *strong labels*) [17]. Recent deep learning models can tag a recording with the set of species present (*weak labels*) [21, 15, 22, 9], but do not predict per-vocalization onset and offsets (*temporal localization*). While sound event detection (SED) models exist that are capable of predicting strong labels [32], in bioacoustics they have focused on detecting fewer than 100 different species (e.g. [20, 7]) due to the high effort required to produce strongly-labeled training data [26]. Despite recent progress in obtaining crowd-sourced strong labels [41], there is still no publicly available collection of strongly labeled data that is sufficiently large and diverse to train a SED system for thousands of species.

In this work, we focus on detecting bird vocalizations, as as birds are highly diverse in species, signal structure, and function [28], while also being well represented in existing training and evaluation data. We present the Bird Communication Detector (BirdCODE), a sound event detection model trained to detect the precise onsets and offsets of vocalizations produced by 9258 species of birds, enabling the study of bird communication at a global scale. We address limitations in training data using three methods: utilizing synthetic strongly-labeled data [10, 42], multi-stage training with strongly pseudolabeled data [38], and a temporal localization-focused training objective for weakly labeled data [40]. We evaluate BirdCODE on 69 strongly-labeled datasets, representing over 1100 bird species, achieving top performance on almost all of these compared to baseline systems. Moreover, despite being designed for sound event detection, BirdCODE achieves state-of-the-art performance on the competitive BirdSet [29] bird sound classification benchmark.

As a first application of BirdCODE, we detected 11.3 million bird vocalizations across 1.3 million recordings in the publicly available citizen-science datasets xeno-canto [43] and iNaturalist [12]. We demonstrate through case studies how these detections can be used to investigate patterns in species-level acoustic communication across the phylogenetic tree of birds, to describe geographic and temporal variation in communication, and to identify potential instances of cross-species communication. Model code and weights are available, as well as detections across these citizen-science datasets.

## Results

### Overview of BirdCODE

We developed BirdCODE, a supervised deep learning model, to solve the following sound event detection (SED) task: given an audio recording containing an unknown (possibly zero) number of bird vocalizations, predict strong labels for these vocalizations, i.e. predict their onset times, offset times, and species identities (Figure 1A). We focused on a zero-shot setting, by which we mean that the model detects bird vocalizations within its training ontology (9258 total) without fine-tuning on the datasets of interest [20] or prompting [10]. BirdCODE predicts, for each frame (frame rate = 7.6 Hz) and each bird species, the probability that that species is vocalizing in that frame. Consecutive frames above a threshold for a species are merged to produce event onsets and offsets.

**Figure 1.**
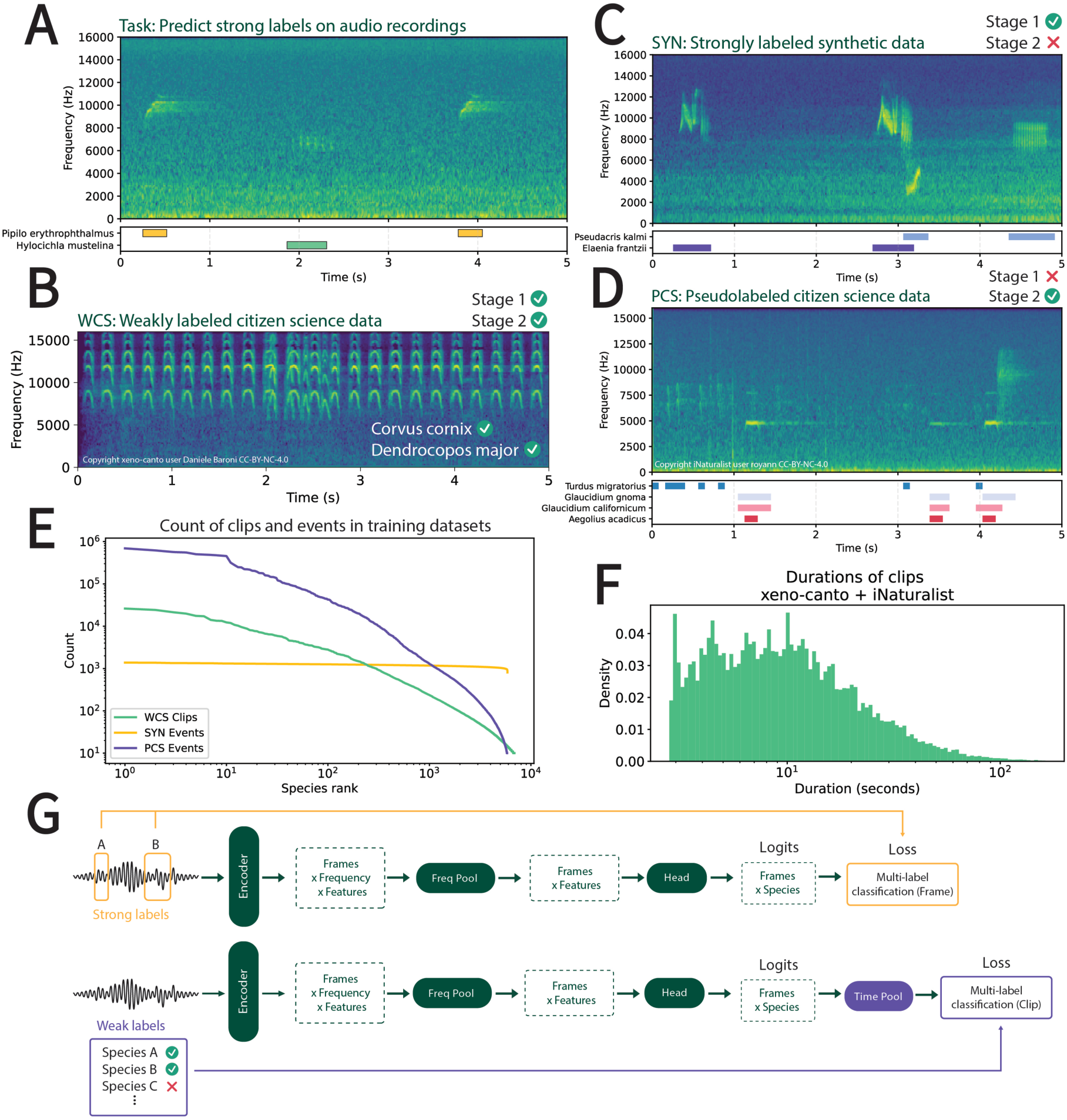
Overview of model development. A) BirdCODE is trained to predict the onset time, offset time, and species (strong labels) of each bird vocalization in an audio recording. B) Large bird sound datasets generated by citizen scientists (xeno-canto and iNaturalist) include tags of one (iNaturalist) or multiple (xeno-canto) species present in the recording. These weakly labeled data (WCS) are used in both stages of the two-stage training pipeline of BirdCODE. C) In the first stage of training, we also use synthetic strongly labeled audio (SYN) which is constructed by pasting short events into longer background recordings, trading realism for temporal precision. D) In the second stage of training, we train on citizen science data with model-predicted strong labels (PCS). E) Distribution of clips and events across species, in the three datasets. The citizen science data (WCS and PCS) has a long-tailed distribution of species, whereas SYN is well-balanced. F) Distribution of durations across WCS/PCS; clips range from a few seconds to several minutes. G) Overview of model training. Strongly labeled data (SYN and PCS) are converted to per-frame species labels (7.6 Hz), which serve as the prediction target. On WCS, the model’s per-frame predictions are pooled into a single per-clip prediction for each species.

Two challenges presented themselves: First, there are roughly 10,000 bird species, meaning that an abundance of training data are required to cover the classes of interest; second, there is a low amount of publicly available strongly labeled data available to use for model training. The data we used to train BirdCODE came from the two large collections of recordings made by citizen scientists, xeno-canto [43] (5.9 *×* 10^5^ recordings, 9237 species) and iNaturalist [12] (6.9*×*10^5^ recordings, 5352 species) (Figure 1B,E,F). While this provided good coverage of species, the recordings were not strongly labeled.

We investigated three strategies for providing a training signal for detecting sound event onsets and offsets. First, we made use of the weakly-labeled citizen science data (henceforth, WCS) by using a custom loss function [40] designed to encourage temporal localization from weakly labeled data (Figure 1G). Second, we generated strongly labeled synthetic acoustic scenes (1 million scenes, 10 seconds each) by pasting short events with known species labels into bird-free background sound recordings [5] (Figure 1C,E; henceforth, SYN). We mixed SYN with WCS during training, using frame-based loss on SYN batches. Finally, following prior work [38], after initial models were trained on WCS and SYN as part of a random hyperparameter search, we ensembled the predictions of six of these across all of the citizen science recordings to obtain predicted strong labels (*pseudolabels*). This dataset (henceforth, PCS), which had the same underlying audio as WCS, replaced SYN in a second stage of training (Figure 1D,E). We used the same initialization (from AudioProtoPNet-20, see below) for both stages; stage 1 was necessary simply to produce pseudolabels for PCS for use in stage 2.

Additionally, we modified and fine-tuned a pre-existing bird classification model AudioProtoPNet-20 (APN-20) [9], which enabled BirdCODE to take advantage of the representation learned in APN-20’s prior training to perform multi-label bird species classification. Specifically, we discarded the temporal pooling in the model head and fine-tuned all model weights, including prototypes. Dropping the pooling layer allowed the model to make high frame-rate multi-label species classifications (7.6 Hz), while simultaneously retaining some of the classification performance learned by the original model.

### Benchmarking BirdCODE’s detection performance

We evaluated the performance of BirdCODE for the zero-shot bird SED task, in which a model detects bird vocalizations without fine-tuning or prompting for the specific evaluation dataset. The main evaluation collection was WABAD [27], which consists of soundscape recordings made with passive acoustic monitors at 72 recording sites. We restricted to the 68 of these recording site datasets which included expert-generated annotations of strong labels (Figure 2A, 4262 recordings total, 1112 species total, 10 min / 111 max / 31.4 average species per dataset). The auxiliary evaluation dataset XCSL, introduced in [14], consists of strongly-labeled focal recordings drawn from xeno-canto (Figure 2A, 967 recordings, 22 species) and held out from training. We used mean average precision (mAP), a class-averaged, decision threshold-free metric, to quantify performance. We report three variants: frame-based mAP, which evaluates predictions by dividing audio into 0.01-second bins and comparing annotations within each bin; and two event-based mAP variants that match predicted events to ground-truth events at an intersection-over-union (IoU) threshold, at tolerances of IoU= 0.5 (mAP@0.5) and IoU= 0.2 (mAP@0.2). As supplemental metrics, we report macro-averaged precision, recall, and F1 at detection threshold of 0.5. Since there was, to our knowledge, no prior large-scale bird SED model, we compared BirdCODE to three baseline systems based on current top-performing multi-label bird species classification models: Perch 2 [21], APN-20 [9], and sl-BEATs-all [22]. We excluded a fourth classification model, BirdNet [15], because its training dataset is not publicly documented, meaning that some or all of the data we used for evaluation may be part of its training set. For each of these classification models, detections were produced by sliding the model across a recording (window size and hop size optimized at 2 sec). This sliding window method reflects typical usage of these classifiers for sound event detection [17].

**Figure 2.**
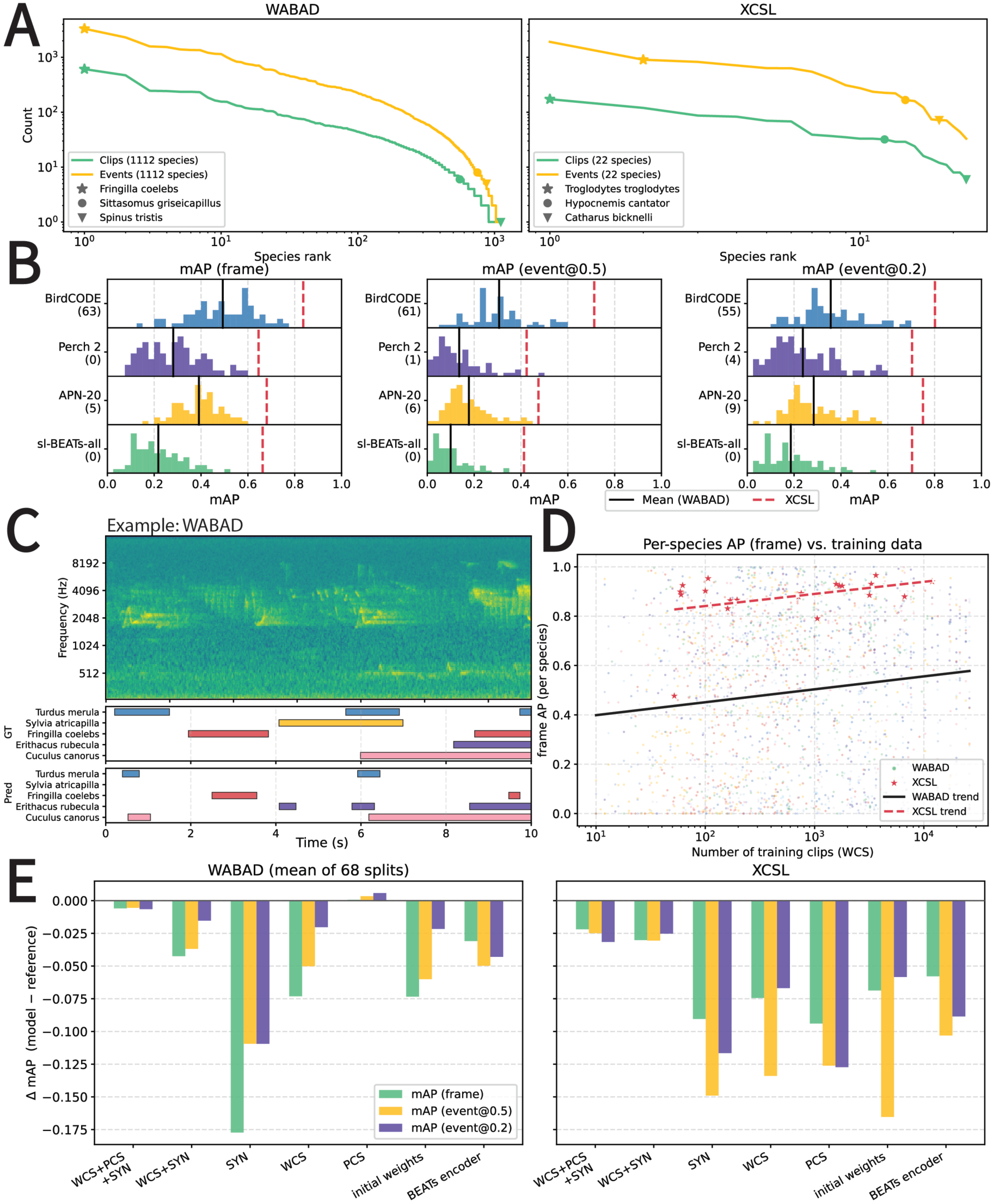
Benchmarking detection performance. A) Species count by rank for our evaluation datasets: WABAD, a collection of strongly-labeled soundscapes recorded at 68 locations, and XCSL, a set of strongly-labeled recordings from xeno-canto. Species with maximum, median, and minimum representation by number of clips in each dataset are marked. B) Performance comparison for BirdCODE against baseline methods using sliding-window classifiers. The number in parentheses below each model name is the number of WABAD datasets where the model achieved the top score in that metric. Histogram of performance on WABAD datasets, with lines for WABAD mean and XCSL. C) Qualitative example of BirdCODE predictions on a clip from WABAD. Ground truth labels made by a human are on top, and predictions by BirdCODE are on the bottom. D) Comparison of per-species performance (frame AP) versus amount of training data, measured as the number of clips in WCS tagged with that species. Trend lines are simple linear regression. Different colors indicate different WABAD datasets. E) Ablation of BirdCODE with different data mixes, using weights of APN-20 without detection-specific fine-tuning, and using a different pretrained encoder (BEATs). Y axis reference: BirdCODE performance.

On WABAD, BirdCODE obtained the highest frame mAP on 63 of 68 datasets, with mean frame mAP of 0.493, an increase of 0.102 above the next highest (Figures 2B and S1; Table S1). BirdCODE performed nearly as well on event-based metrics (mAP@0.5: top on 61, mean score: 0.307, mean increase of 0.123. mAP0.2: top on 55, mean score of 0.357, mean increase of 0.073, Tables S2-S3). On XCSL, BirdCODE obtained the highest score in each metric relative to the baselines (mAP of 0.836, 0.713, 0.802 with increase of 0.156, 0.239, 0.051 on frame, event@0.5, event@0.2, respectively, Tables S1-S3). Of the two event-based metrics, mAP@0.5 penalizes incorrect temporal localization more than mAP@0.2; therefore, the larger difference between BirdCODE and the baselines on mAP@0.5 indicates that at least part of the improved performance is due to better localization than the baseline systems. High performance of BirdCODE relative to baselines continued on supplementary metrics of precision, recall, and F1 (Tables S4-S12). At the detection threshold of 0.5, BirdCODE had relatively high precision (WABAD, frame: 0.86; XCSL, frame: 0.91) but lower recall, especially on the soundscape datasets of WABAD (WABAD, frame: 0.27; XCSL, frame: 0.70). Qualitatively, BirdCODE was able to detect overlapping sounds from multiple species in complex acoustic scenes, although performance was uneven across species (Figure 2C,D). Together, these results demonstrate that BirdCODE is the best overall choice for the zero-shot bird SED task.

We built a statistical model to predict the per-species performance of BirdCODE based on the amount of per-species training data (Figure 2D). A tenfold increase in training clips is associated with a 0.059 increase in per-species frame AP (linear mixed effects model, *p <* 0.001), although variance was high. Recording type (focal vs. soundscape) was included as a fixed effect, and species detected in focal recordings (i.e. XCSL) had AP 0.386 higher than soundscape recordings (i.e. WABAD) on average (*p* = 0.004). The underlying dataset was included as a random effect, and had a modest effect on model performance (69 datasets, interclass correlation coefficient = 0.150), possibly due to differences in recording conditions at different sites.

To isolate what contributed to the high performance of BirdCODE, we conducted a series of ablations (Figure 2E). Of the data mixes available during the first stage of training (WCS only, SYN only, and WCS plus SYN), the combination of WCS and SYN achieved the highest average performance on both datasets, on all metrics. This supports our usage of models trained on this data mix for producing the pseudolabels used in the second stage of training. The final BirdCODE was trained using WCS and PCS; adding SYN diminished average performance on both datasets, on all metrics. On the other hand, training on PCS alone slightly improved performance on WABAD, but substantially diminished performance on XCSL. We also investigated the impact of weight initialization and fine tuning: initializing BirdCODE with random weights entirely failed to train (results not displayed). Including the architectural modifications to APN-20 but not fine-tuning the model led to reduced scores on both datasets, on all metrics. Finally, replacing the AudioProtoPNet encoder with a common alternative [32] (this consists of BEATs [6] plus a randomly initialized linear head) also led to reduced performance on both WABAD and XCSL, for all metrics.

### Benchmarking BirdCODE’s classification performance

BirdCODE can also be used as a multi-label bird species classifier, by applying temporal pooling to the frame-based predictions as during training (Figure 1G). We evaluated the zero-shot bird species classification capabilities of BirdCODE in comparison to Perch 2 [21], APN-20 [9], and sl-BEATs-all [22]. As an evaluation set, we used BirdSet [29], which consists of 7 datasets of soundscape recordings (317732 5-sec recordings total, 402 species total, 65.3 average species / dataset). Following [29], we report class mean average precision (cmAP), area under ROC curve (AUROC), and top-1 accuracy.

Out of the 7 datasets, BirdCODE obtained the top score on 6, 5, and 7 datasets for cmAP, AUROC, and T1 metrics, respectively. (Figure 3A). On average, BirdCODE changed relative to the next best score by 0.059, *−*0.003, and 0.078 on each metric. Together, these results establish BirdCODE as a state-of-the-art model for zero-shot bird sound classification.

**Figure 3.**
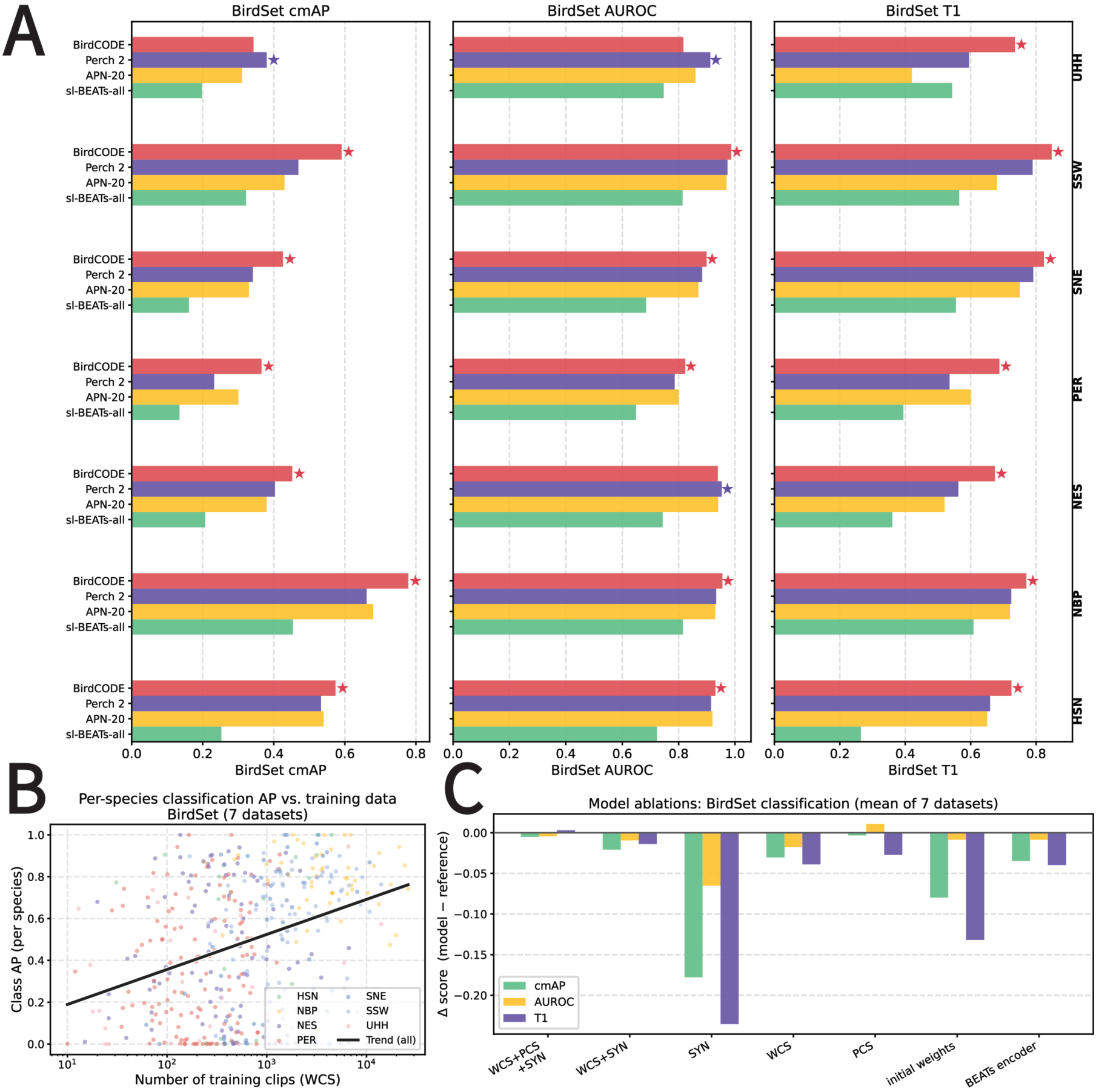
Benchmarking classification performance. A) Quantitative results across BirdSet benchmark, for cMAP, AUROC, and top-1 accuracy metrics, across the 7 evaluation datasets in BirdSet. Stars indicate which model performed best on each dataset and metric. B) Comparison of per-species performance (AP) versus amount of training data, measured as the number of WCS clips tagged with that species. Trend line is simple linear regression. C) Ablation of BirdCODE with different data mixes, using weights of APN-20 without detection-specific fine-tuning, as well as using a different pretrained encoder (BEATs). Y axis reference: BirdCODE performance.

We built a statistical model to predict per-species average precision (AP) based on the amount of available weakly-labeled training data and the underlying evaluation dataset (Figure 3B). A tenfold increase in training data was associated with a 0.103 increase in per-species classification AP (linear mixed effects model, *p<* 0.001), roughly twice that of the SED task. The underlying dataset was included as a random effect, and it had a lower contribution to the total variance than for the SED task (interclass correlation coefficient = 0.110).

We conducted ablations on the data mix used for training (Figure 3C). Of the three data mixes available during the first stage of training (WCS only, SYN only, and WCS plus SYN), the combination of WCS and SYN achieved the highest average performance, on all metrics, again justifying usage of this data mix during the first stage of training. The model trained on a combination of WCS, SYN, and PCS performed essentially the same, on average, as BirdCODE. The model trained on PCS alone improved slightly in terms of AUROC (+0.011), but had lower top-1 accuracy (-0.027); cmAP remained essentially the same as BirdCODE(-0.003). The initial weights model (which is simply APN-20) and the BEATs-based model both obtained lower scores on all three metrics.

### Applications of BirdCODE to animal communication

Using BirdCODE, we detected vocalizations across xeno-canto (587609 recordings, 8581786 detections) and iNaturalist (691761 recordings, 2706044 detections) datasets (Figure 4A, Figures S2-S6). To reduce the number of false positive detections, we cross-referenced recording location data with published species range maps [13], and kept only in-range detections. Performance on these datasets was already validated through benchmarking on XCSL (Figure 2), in which BirdCODE had high precision (frame: 0.91, event@0.5: 0.79, event@0.2: 0.87) and recall (frame: 0.70, event@0.5: 0.77, event@0.2: 0.84) at the detection threshold of 0.5 which was used here. Using these detections, we conducted proof-of-concept studies of bird vocal behavior on four themes: species-level variation in vocal behavior, geographic variation in vocal behavior, cross-species vocal interactions, and temporal variation in vocal behavior.

**Figure 4.**
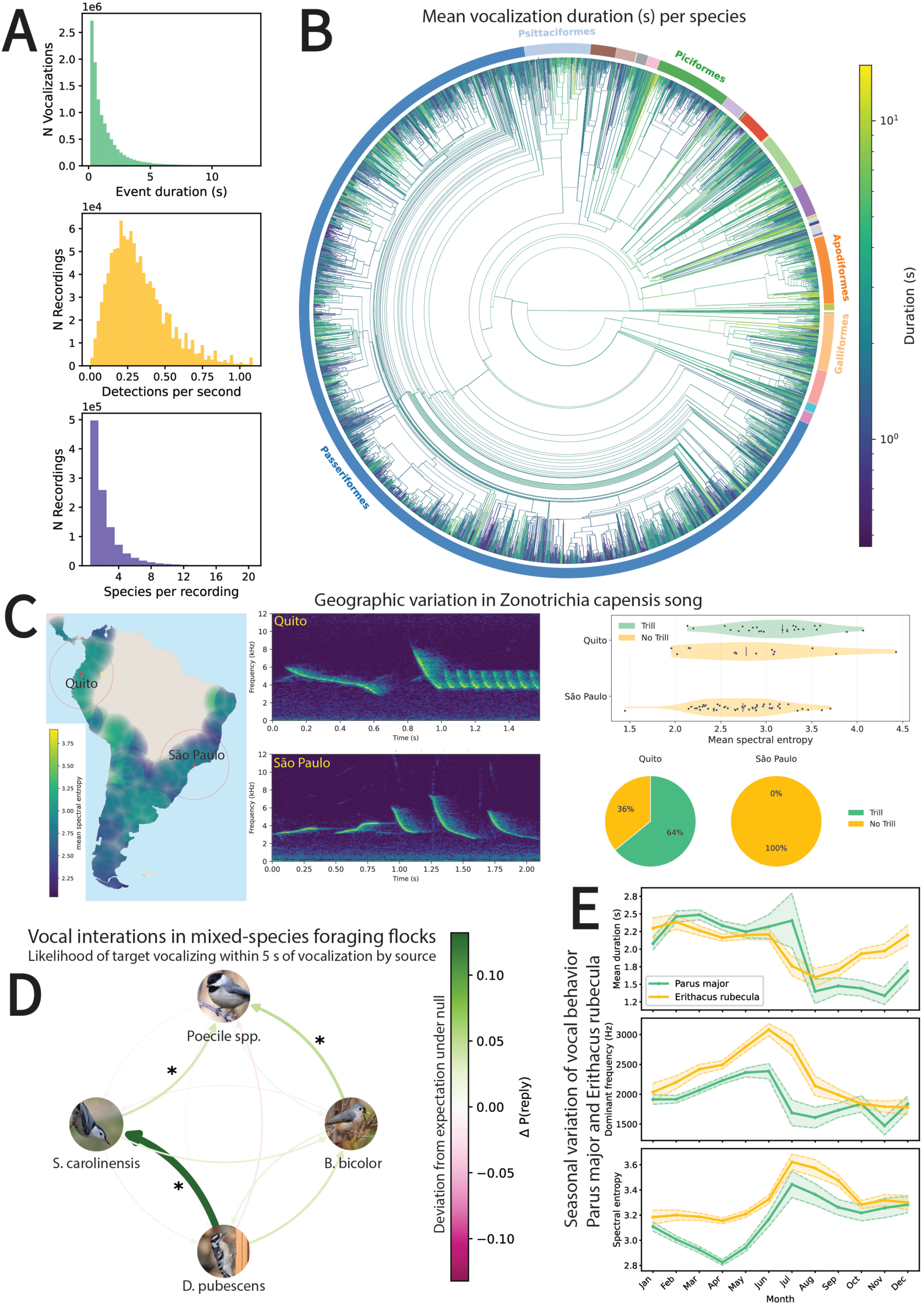
Applications of BirdCODE. A) Summary of detections: duration per detection, rate of vocalization detections in a recording, and number of species detected per recording. B) Phylogenetic tree of birds with *>* 5 recordings containing detections; exterior colors separate different phylogenetic orders, leaf color denotes average duration of vocalizations detected in that species, and interior edge color denotes ancestral state reconstruction. C) Geographic variation in spectral entropy of rufous-collared sparrow song. Left: color denotes average of recordings within 500km radius, and opacity decays with distance from recording location (transparent after 500km). Red circles denote 1000km radius around each city. Center: Spectrograms of songs sampled from near Quito (top shaded region, trill present) and São Paulo (bottom, trill present). Top Right: Spectral entropy of songs with and without trills in each region. Bottom Right: 64% of songs from Quito had trills, whereas none from São Paulo had trills D) Weighted graph of cross-species vocal interactions; edge color and thickness correspond to deviation from null of how strongly vocalizations by one species predict those of the other. Stars indicate significant interactions (Bonferroni-corrected). E) Per-month variation in duration, dominant frequency, and spectral entropy of vocalizations by great tit (*Parus major*) and European robin (*Erithacus rubecula*). Shaded region is 95% confidence interval.

### Application to species-level vocal behavior

Birds produce a diverse set of sounds, and often features of these sounds are more similar in closely related species than distantly related ones [1]. Investigating this typically requires human annotation of individual vocalizations prior to feature extraction; here we instead use the detections produced by BirdCODE. Here, we examined whether there was phylogenetic signal in the average duration of vocalizations recorded in a species (Figure 4B). To decrease label noise, we filtered to detections where the species label matched the recording-level label from the original dataset (4.9 *×* 10^6^ after filtering), and limited to species where there were at least five recordings with detections. For each species, we took the average of detected vocalization durations per-recording, and then took the mean across recordings to obtain per-species durations. To quantify phylogenetic signal, we computed Blomberg’s *K* and Pagel’s *λ* [24] based on the Aves tree from TimeTree.org [18]. For Pagel’s *λ*, we controlled for body size, beak length, and habitat using trait data from AVONET [35]. Blomberg’s K failed to detect a phylogenetic signal (*K* = 1.91 *×* 10*^−^*^8^, permutation test *p* = 0.171), but Pagel’s *λ* detected a strong phylogenetic signal (*λ* = 0.759, likelihood ratio test *p <* 0.001). The divergence between *K* and *λ* is not unprecedented, as *λ* is often a higher-powered test of phylogenetic signal [24].

### Application to geographic variation

We examined geographic variation in song features of the rufous-collared sparrow (*Zonotrichia capensis*), a small songbird widespread in South America (Figure 4D). Songs sometimes include a terminal trill or buzz, which is used by individuals in some areas but not others, possibly due to differences in habitat [36]. We filtered to detections where the species label matched the recording-level label from the original dataset, and plotted a heatmap of the average spectral entropy of vocalizations in a 500-km radius, restricting to vocalizations lasting *>* 0.5 seconds and occurring within October-January. From this, we identified the well-sampled regions around Quito, Ecuador (568 vocalizations, 102 recordings), and São Paulo, Brazil (244 vocalizations, 60 recordings), as containing vocalizations with higher and lower average spectral entropy, respectively. We predicted that trills would have higher average spectral entropy because trills tend to be noisier than non-trills, and that the region with higher spectral entropy would have more trills recorded. We found support for both of these predictions: First, we inspected 50 random vocalizations from the 1000-km radius around each city. Some files (13, 13%) were skipped because they were not songs or were false positive detections (10), or because the bird vocalization was too faint (3). Almost all inspected vocalizations were songs (90%). Trilled songs had higher spectral entropy on average (one-tailed Welch *t* = 2.901, *df* = 47.5, *p* = 0.003). Near São Paulo, none of songs contained trills, whereas near Quito, 25/39 (64%) songs contained trills. Nontrilled songs from the two regions did not have significantly different spectral entropy (one-tailed Welch *t* = 0.683, *df* = 16.0, *p* = 0.505).

### Application to cross-species interactions

Communication occurs across species in a variety of contexts, include predator avoidance, territoriality, and aggression [3]. The timing between vocalizations of different species may provide insight into differing social roles in cross-species interactions [39]. During the winter in North America, mixed-species foraging flocks form which include parids (black-capped chickadee *Poecile atricapillus*, Carolina chickadee *Poecile carolinensis*, and tufted titmouse *Baeolophus bicolor*) as well as other satellite species (white-breasted nuthatch *Sitta carolinensis*, downy woodpecker *Dryobates pubescens*, and others) [23]. We analyzed recordings made in September-February which included detections at least two of these five species, treating the chickadees as one species since they fulfill similar ecological roles and have minimally overlapping ranges. For each ordered pair of species (A,B), for each recording, we computed the probability that B would vocalize within the *T* = 5 seconds following a vocalization made by A (Figure 4E). These probabilities were averaged across all recordings containing the pair, to give a single probability per ordered pair. We tested whether this value exceeded the expectation under simulated data in which the timing of one species’ vocalizations was circularly shifted by a random time interval.

We found three species pairs with a significant effect: tufted titmouse vocalizations predict chickadee vocalizations (deviation in probability = 0.047, one-tailed corrected *p* = 0.012), white breasted-nuthatch vocalizations predict chickadee vocalizations (deviation in probability = 0.036, one-tailed corrected *p* = 0.048), and vocalizations or drumming by downy woodpecker predict white-breasted nuthatch vocalizations (deviation in probability from null = 0.132, one-tailed corrected *p* = 0.012). The results for chickadee responses were robust to variation *±*1 sec in T (corrected *p ≤* 0.012 in all cases), although downy woodpecker vocalizations were not predictive of anything when *T* = 4 (*p ≥* 0.780). These results align with studies showing that titmice lead flock movement [8], while chickadees remain closely connected but typically subordinate to titmice [30]. Chickadees’ responses to titmice and nuthatches are consistent with findings showing that chickadees are often first to approach playbacks of heterospecific calls [4]. The nuthatch–chickadee and nuthatch–woodpecker associations also corroborate work showing close social ties between nuthatches and both species in mixed flocks [39]. While correlation in call timing does not imply that one species vocalizing causes another to do so, these results suggest directions for future experiments in this multi-species system.

### Application to seasonal variation

We performed a qualitative investigation of month-by-month variation in acoustic features (duration, dominant frequency, spectral entropy) of the vocalizations of two common European songbirds, the great tit (*Parus major*, 75509 vocalizations, 12757 recordings) and the European robin (*Erithacus rubecula*, 106028 vocalizations, 13025 recordings). For both, we filtered to detections where the species label matched the recording-level label from the original dataset. We plotted the average value of each feature per month (Figure 4C). Great tit vocal features exhibited two qualitative modes: vocalizations were, on average, longer, higher frequency, and lower entropy in the first half of the year, and shorter, lower frequency, and higher entropy during the second half. European robin vocalizations also exhibited two modes, with vocalizations made from July-Sept being on average longer and higher entropy than other months. Dominant frequency was elevated during a slightly earlier period, from May-July. Both of these patterns recover known behaviors. Great tits sing from late winter until the end of nesting in June or July [2]; their songs are high-pitched and tonal, and can be several seconds long, versus calls which are shorter and include lower-pitched noisy chatters. European robins sing year-round, but reduce territorial behavior during the post-reproductive season, between mid-summer and fall [33]. The change in duration and entropy in July-Sept may reflect a decrease in singing relative to other call types; on the other hand, the increase in dominant frequency in the spring and early summer may reflect acoustic differences between the spring and autumn songs [2].

## Discussion

We introduced BirdCODE, a method to effectively detect and classify 9258 bird species in audio recordings without any need for additional training, enabling novel high throughput applications in animal communication. In recent years, advances in biodiversity monitoring [19], wildlife management [11], and movement ecology [25] have been driven by methods for collecting, curating, and interpreting very large datasets; simlarly, we expect BirdCODE to contribute to the emerging “big data” era of animal communication research [34].

The central challenge of developing BirdCODE was the lack of strongly labeled training data covering thousands of species, and we used three strategies for overcoming this challenge: training with localization-focused loss on weakly labeled data (WCS), training with strongly labeled synthetic data (SYN), and training with strongly pseudolabeled data (PCS). Using SYN, on its own, led to low performance, likely due to lack of realism in the synthetic scenes. However, the combination of SYN with WCS outperformed the model trained on WCS alone, suggesting that some amount of strongly labeled data, even if lacking in realism, can boost detection performance. The model trained on PCS alone improved in some cases on BirdCODE, which was trained with a combination of PCS and WCS. One explanation is that in PCS, background species were detected that were missing from original weak labels, as pseudolabeling has been used to address specifically this problem in bird sound classification [38]. This may also explain why BirdCODE outperformed the model trained only on WCS for the classification task. The notable exception to this were the results on XCSL, where model trained on PCS alone underperformed BirdCODE. The XCSL dataset was constructed in [14] specifically to focus on similar-sounding pairs of birds. Therefore, a likely explanation for the difference in performance between these two models is that noise in the pseudolabels of PCS introduces confusions between similar sounding species, which training on WCS partially corrects.

An inherent limitation in bioacoustic SED is that the basic unit, a vocalization, may differ between datasets depending on annotation protocols for merging or splitting sounds that appear near each other [17]. In postprocessing, we merged detections that were *<* 1 second apart, but this choice was arbitrary and not optimized. Optimal postprocessing choices may vary between datasets, and even between species within a dataset. A limitation of BirdCODE specifically is that it detects only birds, as opposed to a wider set of taxa, and it classifies vocalizations to species, as opposed to, for example, vocalization types within a species. While both of these limitations could be addressed with additional training data (which does exist, to some extent, on platforms like xeno-canto and iNaturalist), there would remain a lack of strongly-labeled data for evaluation of model capabilities beyond bird species detection. Developing a strongly labeled dataset with wide geographic and taxonomic coverage (analogous to WABAD) would be a prerequisite for developing these extensions to BirdCODE. We did not consider alternate sound event detection frameworks, such as those based on onset detection [20] or open set detection [42]; the benefits and drawbacks of each of these approaches for large-scale detection of animal sounds merits further study.

Applications using BirdCODE on citizen science data must carefully consider how to correct for confounds in the data, such as variation in background noise correlated with geography or recording time. They also must address sampling biases, such as under representation of the Global South, and over representation of behaviors which may be considered more interesting to a recordist (e.g., singing) [37]. While we performed some data filtering in our proof-of-concept applications, further development of filtering methods will help increase confidence in conclusions. Similarly, validating outputs, as in our investigation of geographic variation, will be essential. We focused applications on citizen science data, but we also demonstrated through evaluation on WABAD that BirdCODE can be successfully applied to soundscape data collected by passive acoustic recorders. Applications to data collected by large deployments of passive acoustic sensors, as in [16], could provide another avenue towards uncovering patterns in animal communication. We also expect the detections produced by BirdCODE can be used in other areas of computational bioacoustics, such as developing sequence models of animal communication, and as a data source for future bioacoustics foundation models.

## Data and Code availability

Trained model weights, evaluation code, inference API, and precomputed detections are available at https://github.com/earthspecies/sound-event-detection. Code to produce figures used in this paper is available at https://zenodo.org/records/21711348.

## Contributions

Conceptualization: AF, BH, DR, MM, EC, MC, LSJ, SK, IN, MG Methodology: AF, BH, DR, MM, MG Software: AF, BH, DR, MM, MA, EM, GN Validation: AF, BH, DR, MM Formal analysis: AF, BH, DR Investigation: AF, BH, DR Resources: MA, EM, GN Data curation: AF, BH, DR, MA, EM, GN Writing – original draft: AF, BH Writing – review & editing: AF, BH, EC, MC, LSJ, SK, IN, MG Visualization: AF, BH, EC, MC, LSJ, IN Supervision: BH, DR, MM, EC, MG Project administration: BH, DR Funding acquisition: EC, SK, MG

## Acknowledgments

We thank Ellen Gilsenan-McMahon for project administration. We are grateful to Laura Hay, Chiara Semenzin, Gregory Yauney, Emmanuel Fernandez, and Raphael Korach for participating in discussions regarding this work. We thank Diane Kim for assistance with preparing figures. We thank Olivier Pietquin for general supervision of the research group. Finally, we thank the broader Earth Species Project team for their support.

## 1 Online Methods

### 1.1 Dataset

#### 1.1.1 Ontology

Since we combined multiple models and datasets, each of which had its own naming conventions, it was necessary to convert scientific names to a common format. Throughout, species names were resolved to “canonical” names under the Global Biodiversity Information Facility (GBIF) 2021 backbone taxonomy [6], resolving synonyms to their accepted usage.

#### 1.1.2 Training set

##### WCS: Weakly labeled citizen science data

The model was trained on two weakly labeled citizen science datasets of animal sound recordings: iNaturalist [9] and Xeno-canto [31]. We used recordings downloaded on February 3, 2026 and excluded all recordings with a no derivatives license, as well as recordings present in the XCSL evaluation dataset. The iNaturalist dataset comprises 699,217 recordings totaling 3,772 hours, each labeled with a single focal species. The Xeno-canto dataset comprises 587,628 recordings totaling 8,215 hours; in addition to a focal species, each recording may include a list of background species derived from the recordistannotated “associated taxa” field. There were 9258 bird species represented, and 6540 species with at least 10 recordings. Recordings from non-birds were included, but non-bird labels were dropped. To reduce long-tail imbalance, species with fewer than 75 recordings were upsampled by repeating recordings until each species reached 75 recordings or each recording had been repeated 10 times, whichever came first.

##### SYN: Strongly labeled synthetic data

We generated 1 million scenes, each 10-seconds long, by pasting short foreground recordings of bird vocalizations into longer background recordings. The foreground recordings had species labels and were tightly cropped to the sound event, making it possible to generate strong labels on the synthetic scene (i.e. event onsets, offsets, and species labels).

To obtain foreground bird vocalizations, we began with the foreground “pseudovocalization” dataset developed in [8]. This dataset consisted of 5.4 *×* 10^6^ short (typically *<* 2 second) recordings (577 hours), which had been denoised and filtered to remove recordings which did not resemble animal sounds. This dataset did not initially have species labels, therefore we developed a process to automatically assign these labels: Each short foreground recording from [8] was originally cropped from a longer recording in [9, 31] which included a weak species label (the *metadata species*). Using an internal species classification model (an EfficientNet-B0 [28] classifier trained on 8,462 animal species using procedures from [18]), we predicted the probability of each of these 8,462 species occurring in the foreground recording. We retained a foreground recording if the metadata species occurred in the top-25 predicted species, and the sigmoid probability of the metadata species also exceeded 0.05; otherwise, the recording was discarded. Finally, foreground recordings from species with *<* 5 foreground recordings were discarded. After filtering, 601,179 recordings spanning 6,280 species were retained. Retained recordings were labeled with their metadata species and resampled to 32kHz. Since the metadata species was originally ascribed by the human recordist, this filtering process served the purpose of isolating the vocalizations which were made by the metadata species.

Background soundscapes were drawn from the Silent Cities dataset [1], a collection of 4.5 *×* 10^4^ hours of urban soundscape recordings across 317 locations made in 2020. We downloaded 461 tar-archived batches (159 unique monitoring locations) and randomly subsampled up to 500 files per archive. Candidate backgrounds were then passed through two successive filtering stages to remove files containing significant biological activity. In the first stage, the same internal EfficientNet-B0 classifier was applied to each full 10-second file; files for which any animal class prediction exceeded a sigmoid probability of 0.1 were discarded. In the second stage, remaining files were screened with a second classifier (`esp_aves2_effnetb0_bio` from [18]) and files with a maximum per-class softmax probability above 0.2 were removed. Both thresholds were selected to exclude soundscapes containing audible species vocalizations while retaining natural ambient textures (wind, rain, distant traffic). After filtering, 396,662 ten-second clips at 32 kHz were retained, totaling approximately 1,102 hours. Retained files were resampled to 32 kHz and cropped to 10 seconds. Retained files were manually spot-checked and had almost no animal sounds present.

Finally, synthetic soundscapes were constructed by mixing filtered foreground clips onto filtered background recordings. For each scene, the number of focal species was drawn from a truncated Poisson distribution (*λ* = 2.0, support [1, 3]), and for each species the number of vocalization events was drawn independently from a second truncated Poisson (*λ* = 3.5, support [1, 100]). A single background file was selected uniformly at random. Each foreground clip was gain-adjusted to a target level drawn uniformly from [*−*18, +6] dB relative to the RMS of the background. Events were placed at uniformly random onset times within the 10-second scene, with overlaps between events permitted. The mixed waveform was clipped to [*−*1, 1] and saved as a 16-bit WAV at 32 kHz. Annotations were stored with the start time, stop time, and species label of each event; overlapping events from the same species were merged. In total, 1 million unique scenes were generated. Upon manual inspection, audio artifacts from the generation process were rare; however, the intervals between vocalizations and the exact species which co-occurred in scenes were often non-naturalistic.

##### PCS: Strongly pseudolabeled citizen science data

We produced pseudolabels (i.e. labels predicted by a model) on the audio from xeno-canto and iNaturalist. The resulting dataset PCS therefore had the same audio as in WCS, but a different set of annotations. To do so, we used six detection models which were trained independently on the combination of WCS and SYN data (i.e. during stage 1). These models were trained during initial sweeps of loss and data augmentation hyperparameters, and selected based on their performance on the validation datasets.

For each audio clip, the frame-level probability arrays from all six models were averaged to produce a single ensembled probability array. The ensembled frame probabilities were converted to event-level detections using a fixed probability threshold of 0.4. Any frame exceeding this threshold was retained as a candidate detection. This probability threshold was selected from *{*0.4, 0.5*}* based on validation set performance of the stage 2 model (i.e. a threshold of 0.4 led to better performance than 0.5). These frame-level prediction ensembles were not postprocessed in any way, and they were taken as the annotations for PCS.

#### 1.1.3 Evaluation data

##### Detection

We evaluate detection performance on the World Annotated Bird Acoustic Dataset (WABAD) [23], a collection of strongly annotated soundscape recordings, and XCSL [11], a collection of strongly annotated recordings from xeno-canto (Table 1). The original WABAD dataset consists of recordings from 72 sites; we excluded four sites which included only weak labels. For validation (i.e. hyperparameter tuning and model selection), we used the Powdermill dataset [3], a collection of strongly annotated soundscape recordings from Pennsylvania, USA. All annotation labels were standardized to GBIF canonical names using the same procedure as the training data. When evaluating on any individual dataset or recording site, model predictions were filtered to the species annotated in that dataset before computing metrics (see Section 1.7).

**Table 1.**
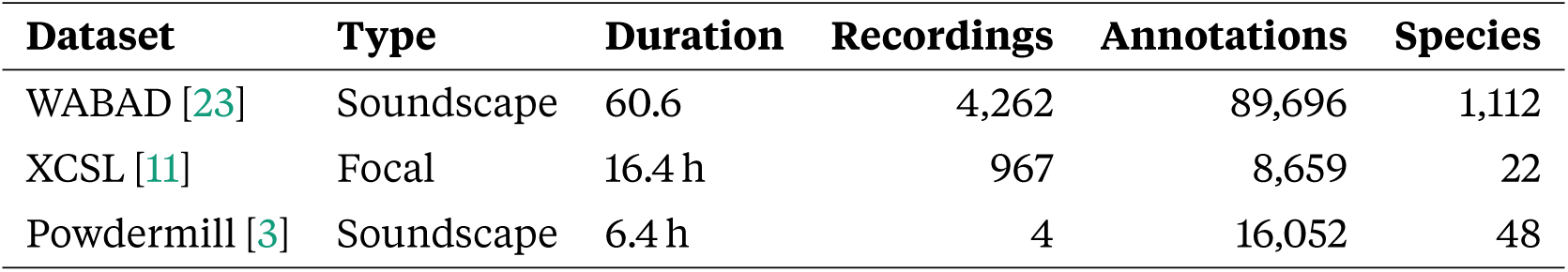
Detection evaluation datasets. Species counts reflect unique annotated taxa after GBIF standardization.

##### Classification

For evaluating clip-level classification we use the test set of BirdSet [24]. BirdSet consists of seven datasets of soundscape recordings, each a collection of 5-s clips annotated with the set of species that occur in the clip. Following [17] we do not use the eighth POW dataset for evaluation; note also this dataset was derived from the Powdermill dataset we had already used as a detection validation set. In contrast to [24], and following [17], we do not use the train split of BirdSet and only use the test set to measure zero-shot classification performance.

### 1.2 Model Architecture

Our training method assumed a model which maps an audio waveform *x ∈* ℝ^T^ (*T* is the number of samples) to an array of probabilities *ỹ∈* ℝ^F^ *^×^*^C^ (*F* is the number of frames, *C* is the number of species). The input sample rate and output frame rate are assumed constant across audio clips. The value *ỹ*_f,c_ at a given frame *f*, for a given species *c*, represents the model’s prediction of the conditional probability P(*c* is making a sound at *f | x*). In practice, we used a modified APN-20 [7] architecture. We additionally compared this to a model based on a BEATs encoder. Both are described in detail below.

### 1.3 Training Objective

During training, weakly labeled data and strongly labeled data are passed to the model in separate batches. For strongly labeled data, focal loss [13] is applied per-frame to the logits. Let *y ∈* {0, 1}^F^ *^×^*^C^ be per-frame ground truth labels, where *y*_f,c_ = 1 if species *c* is making a sound at *f* and 0 otherwise. Then, letting *ỹ* be as above the loss is

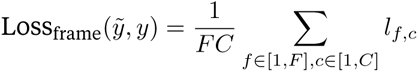

where

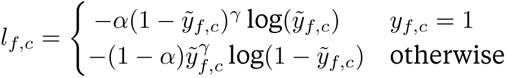

and *α ∈* [0, 1] and *γ ≥* 0 are hyperparameters.

For weakly labeled data, frame-level predictions are pooled in time to a clip-level prediction *ỹ*_clip_ *∈* ℝ^C^ using the “softmax” pooling from [30]:

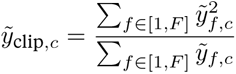

These predictions compared to clip-level binary labels *y*_clip_ *∈* {0, 1}^C^ via a focal loss:

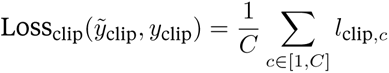

where

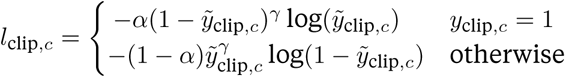

and *α, γ* are as before. The total loss is the sum of the batch-averaged clip-level and frame-level losses.

### 1.4 Data Augmentations

Three augmentations are applied at the batch level during training.

#### Additive audio mixing

With some probability (*p* = 0.56 per round, up to 3 rounds), each training clip is mixed with one another clip sampled from the same batch, and labels are unioned (clip- or frame-wise maximum).

Because the majority of weakly labeled training data consists of focal recordings containing a single species, this augmentation introduces soundscape-like training examples with overlapping vocalizations from multiple species.

#### Noise injection

Noise augmentation used the set of Silentcities recordings that were used as background recordings in SYN. These recordings were used in two ways: with some probability (*p* = 0.45), a noise clip is added to the training audio at random gain (uniform in [*−*5, 0] dB), leaving labels unchanged; independently, with a smaller probability (*p* = 0.02), the training audio is replaced entirely by a noise clip and all labels are zeroed, creating explicit negative examples.

#### Random gain

A random gain, sampled uniformly in dB (clip-level: [*−*5, 0] dB; frame-level: [*−*1, 0] dB), is applied to each training clip. The smaller range in frame-level data was adopted to reduce instances where quiet background vocalizations with annotations were rendered inaudible.

### 1.5 Implementation Details

#### AudioProtoPNet encoder

We modify and fine-tune the pre-trained AudioProtoPNet-20 (APN-20) [7] model, which was originally designed to predict which of 9736 bird species occur in an audio clip (i.e. predict weak labels). Audio (sampling rate: 32kHz) is converted to a log-mel spectrogram, then passed through a ConvNeXt-B [14] backbone to obtain frame-based feature vectors. The model computes cosine similarities between learned prototypes and a ConvNeXt feature map at each spatial position, then applies a global max-pool over both frequency and time to produce a clip-level prototype similarity measurements. Finally, these similarities are passed through a linear head, which uses non-negative weights connecting each class only to its own prototype similarities [7]. In our modification, we remove the max-pooling over time and retain only the frequency-axis reduction. These per-frame similarities are then passed through the linear prediction head, independently at each time step. Therefore, the model natively produces per-frame predictions for each species. After converting the original Audio-ProtoPNet taxonomy labels to GBIF canonical names and deduplicating synonyms, the model covers 9422 species at 7.6 Hz (38 frames per 5-second window). This pre-trained model is then fine-tuned using our training methodology. In the data used for training BirdCODE (i.e. PCS and WCS), there were 9258 bird species represented, and 6540 with *≥* 10 recordings in the WCS dataset. Therefore, while BirdCODE produces predictions for 9422 species, only 9258 were specifically fine tuned for the purpose of detection. The number of training clips per species is available through the provided code.

#### Training procedure

Models were implemented and in Pytorch [22]. Training took place on a cluster with four H100 GPUs. All hyperparameters were tuned in initial experiments based on performance on the Powdermill validation set. The ConvNeXt backbone was frozen for an initial warmup period (0.25 epochs) then unfrozen for end-to-end fine-tuning with a second learning rate warmup period (also 0.25 epochs). We use AdamW [15] with a learning rate of 6 *×* 10*^−^*^4^, weight decay 0.01, linear warmup and cosine annealing with bfloat16 mixed precision. Training runs for 20 epochs, with a batch size of 32 and window size of 5 seconds. Audio recordings and strong labels were randomly cropped to 5 seconds at each epoch, leaving weak labels unmodified. The checkpoint with the best frame-level mAP on Powdermill was selected for evaluation.

Other hyperparameters are as follows: Focal loss: *α* = 0.25, *γ* = 2, label smoothing = 0.1. Additive audio mixing: probability 0.56, up to 4 sources. Noise injection: mix probability 0.45, gain range [*−*5, 0] dB; replace probability 0.02. Random gain: [*−*5, 0] dB for weakly labeled data, [*−*1, 0] dB for strongly labeled data.

### 1.6 Inference and postprocessing

During inference, BirdCODE makes predictions on windowed audio files (window size = 5 sec, hop size = 2.5 sec), keeping only predictions in the center frames of each window to reduce edge artifacts. Frame-level probabilities are converted to discrete detection events by applying a confidence threshold, then four postprocessing steps are applied sequentially. First, consecutive frames with detections of the same species are merged into events, and assigned an event probability score equal to the average probability across the frames. Second, events of the same species separated by less than a maximum gap (length = 1.0 seconds) are merged into a single event; the score of merged events is taken to be the maximum across the events. Next, predictions are filtered: for evaluation, we filtered all model predictions to remove species not present a dataset, and for applications, we filtered predictions based on published species range estimates (described below). Finally, non-maximal suppression [19] removes overlapping events across species at an IoU threshold of 0.8, greedily preferring to keep events with a higher probability score.

### 1.7 Detection performance metrics

To measure detection performance, we report two variants of mean average precision (mAP): frame and event mAP. When evaluating on a given dataset, model predictions are filtered to only the species annotated in that dataset before computing metrics. For each, we sweep 101 detection thresholds linearly spaced from 0.0 to 1.0 to form a precision-recall curve for each species.

#### Frame mAP

At each threshold, per-frame probabilities are binarized and compared to perframe ground truth annotations. Predictions and annotations are both resampled to 100Hz, independent of the model being evaluated. For each threshold and class, frame-level true positives, false positives, and false negatives are counted; a precision-recall curve is computed and interpolated following the Pascal VOC protocol [4], and the area under the curve gives per-class average precision. Frame mAP is the unweighted mean over all classes present in the ground truth.

#### Event mAP

At each threshold, per-frame probabilities are postprocessed into events, as described above. Predicted and ground-truth events are matched based on temporal intersection-over-union (IoU) rather than compared per-frame. The matching procedure follows [20, 8, 16]: a maximum-cardinality bipartite matching is computed between predicted and ground-truth events; a matched pair counts as a true positive if its IoU exceeds a minimum threshold. We report event mAP at IoU thresholds of 0.2 and 0.5. Per-class average precision and macro-averaging follow the same procedure as frame mAP.

#### Supplementary metrics

In addition to mAP, we report precision (i.e. the ratio of true positive detections to the total number of detections), recall (i.e. the ratio of true positive detections to the total number of ground-truth annotations), and F1 (the harmonic mean of precision and recall). These metrics are not threshold-free and are computed at a detection threshold of 0.5, which is the same threshold we employed for all applications. Scores are computed per-species and then averaged. As with mAP, we report frame-based and event-based variants.

### 1.8 Baselines and Comparison Methods

#### Sliding window classifiers

A pretrained audio classifier can serve as a simple detector by sliding it over a recording. At each step, a short analysis window of audio is extracted, zero-padded to the classifier’s expected input length, and classified; the hop between steps determines the output temporal resolution. Where consecutive windows overlap, their logits are averaged before applying the output activation. We evaluated three such sliding classifiers with a 2-second hop and window size (2 Hz output): Perch 2.0 [17], a supervised EfficientNet-based bird sound classifier (5-second native window, 32 kHz); sl_beats_all [18], a self-supervised audio transformer fine-tuned on bioacoustic classification (10-second native window, 16 kHz); and AudioProtoPNet [7], a prototype-based ConvNeXt model trained on multi-label bird sound classification (5-second native window, 32 kHz). For all models, we compared a 2-second window to the native analysis window (5- or 10-second, depending on the model), and also to a shorter 1-second window and hop size; in all cases the 2-second window led to higher detection performance and we report those scores.

#### AudioProtoPNet without time pooling

As an ablation of our method, we evaluated a version of BirdCODE that adopted the architectural modifications to APN-20 which are described above, but was not fine-tuned.

#### BEATs encoder

As an ablation of our method, we evaluated a detection model based on a BEATs [2] encoder, as this is a common approach for sound event detection tasks [26]. BEATs produces embeddings at 8 frequency patches per time step; these are concatenated to yield a 6,144-dimensional embedding (768 *×* 8) at 12.4 Hz (62 frames per 5-second window). These per-frame embeddings are finally passed through a linear prediction head. We initialize using the `iter3+AS2M` checkpoint for the encoder and used a randomly initialized linear head. While BEATs was pre-trained on audio at 16kHz, we feed 32kHz audio because many bird vocalizations are above the Nyquist frequency of 8kHz and because the BEATs representation has previously been shown useful, even on audio input at nonnative sampling rates [16]. Training hyperparameters are the same as for other models.

#### Statistical analysis

To quantify how detection and classification performance scale with the amount of training data available per species, we fit linear mixed-effects models (Python package `statsmodels` [27]) to per-species average precision (AP) scores. In both models we included the base-10 logarithm of the number of training clips in WCS for each species as a fixed effect. For the detection performance model, we also included a binary indicator distinguishing focal recordings (XCSL) from soundscape recordings (WABAD). For both models, random intercept was included for each evaluation dataset to account for the non-independence of species scores assessed on the same dataset. For the detection analysis (Fig. 2), the response variable was per-species frame-level AP. For the classification analysis (Fig. 3), the response variable was per-species classification AP.

### 1.9 Methods for applications

#### 1.9.1 Inference on xeno-canto and iNaturalist

Using BirdCODE, we detected bird vocalizations on the xeno-canto and iNaturalist audio that was used in WCS and PCS. Detections of species outside their typical range were removed prior to non-maximal suppression. We compared to species range predictions published by iNaturalist [10], removing out-of-range detections. This resulted in 8581786 detected vocalizations for xeno-canto and 2706044 detected vocalizations for iNaturalist.

#### 1.9.2 Phylogenetic analysis of vocal duration

To test whether song duration is phylogenetically conserved across birds, we computed per-species mean detection duration and mapped these values onto a time-calibrated phylogeny of Aves. For each audio clip, we used the BirdCODE detections to identify all detections assigned to a given species. Within each clip we averaged the detection durations; the resulting per-clip means were then averaged across clips to yield a single per-species mean duration. To reduce the influence of misidentified recordings, we applied a canonical filter: only detections whose model-assigned species label matched the clip-level ground-truth species label were retained. Species with fewer than five recordings were excluded. The top and bottom 1% of species-level duration values were trimmed before all downstream analyses to remove outlier estimates driven by very short or very long non-representative detections. We used a time-calibrated supertree of Aves from TimeTree.org [12]. The tree was pruned to the set of species that passed the recording-count and outlier filters, resulting in a 3504-species tree.

Per-species mean duration was log-transformed prior to analysis. Blomberg’s *K* was estimated using the `phylosig` from the R package `phytools` [25] (10,000 permutations) to quantify how much phylogenetic clustering of trait values exceeds the expectation under Brownian motion. To compute Pagel’s *λ* (while simultaneously accounting for body size, beak morphology, and habitat), we fitted a phylogenetic generalised least-squares (PGLS) model using the function `pgls` from the R package `caper` [21]. Species-level trait data (culmen length, body mass, habitat category) were drawn from the AVONET database [29]. Beak (culmen) length and body mass were log-transformed, and habitat categories represented by fewer than 50 species in the analysis sample were collapsed into an “Other” category. Pagel’s *λ* was estimated jointly with the regression coefficients by ML. A second model was fit with *λ* fixed to 10*^−^*^6^, and the two models were compared by a likelihood-ratio test (*χ*^2^ on 1 d.f.).

The full Aves timetree was drawn as a circular phylogram with tip points colored by taxonomic order. Ancestral state estimates for branch coloring were obtained by maximum-likelihood reconstruction under Brownian motion (Felsenstein pruning) [5], and branch segments were rendered as color gradients interpolating the reconstructed values at parent and child nodes.

#### 1.9.3 Seasonal variation in vocal features

For each audio clip in both datasets, we extracted the re cording month; Clips lacking this information were excluded. Vocalization onsets and offsets were identified within each clip based on BirdCODE detections. To reduce the influence of misidentified recordings, we applied a canonical filter: only detections whose model-assigned species label matched the clip-level ground-truth species label were retained. Acoustic features were extracted from 32kHz audio using a short-time Fourier transform (1024 filters, 512-sample hop length), with power spectra high-pass filtered above 50 Hz. Dominant frequency was identified per frame as the frequency bin of maximum power and then averaged across frames. Mean spectral entropy was computed by normalizing each frame’s power spectrum to a probability distribution and averaging the per-frame Shannon entropy across all frames. For each clip *×* species combination, acoustic features were averaged across all detections of that species within that clip, yielding one set of features per clip. Plots reflect per-month mean and 95% confidence intervals

#### 1.9.4 Geographic variation in vocal features

To characterize geographic variation in the vocalizations of Zonotrichia capensis, we collated recordings which 1) included BirdCODE detections of Z. capensis, 2) had clip metadata indicating that Z. capensis was present, 3) had GPS coordinates available, and 4) were made in October– January. For each qualifying recording, we excluded detections of *<* 0.5 seconds in order to focus on songs of Z. capensis, which are usually longer than 1 second, rather than calls. For each detected vocalization, we computed spectral entropy (as in Section 1.9.3) and then computed the per-recording mean. Outliers outside of 2 standard deviations of the mean were removed, as were recordings lacking at least one other sample within 500 km. The resulting records were projected onto a 0.1° geographic grid; each land-surface grid cell was assigned the mean feature value across all samples within 500 km, with alpha transparency scaled by the squared ratio of nearest-sample distance to the 500 km bandwidth, so that sparsely sampled regions fade smoothly. We defined two focal regions centered on Quito, Ecuador and São Paulo, Brazil (radius 1,000 km each), based on observed differences in spectral entropy in these regions. From vocalizations within each region, 50 clips were drawn uniformly at random. Audio waveforms and spectrograms were embedded in an HTML annotation interface. One rater (BH) classified each vocalization as trill, no-trill, or skip. Vocalizations were presented in a random order and the origin location was not visible to the rater.

#### 1.9.5 Cross-species interactions

To quantify directed vocal coordination within a mixed-species community, we computed pairwise reply probabilities among four species groups: Poecile spp. (collapsing P. atricapillus and P. carolinensis), Baeolophus bicolor, Dryobates pubescens, and Sitta carolinensis. Recordings from were filtered to winter months (September–February) and, for each ordered pair of species (focal species, nonfocal species), we restricted to clips in which both species were detected by BirdCODE. For each focal vocalization ending at time *t*, we defined it as eligible if at least *T* = 5 seconds remained in the recording, ensuring the full response window was observable. Reply probability for a given recording was computed as the proportion of eligible focal vocalizations followed by at least one vocalization from the nonfocal species within T seconds; per-recording values were then averaged across all qualifying recordings. To assess statistical significance, we constructed a null distribution via 1,000 circular time-shift permutations: for each permutation, focal and nonfocal vocalizations within every qualifying recording were independently shifted by a random amount drawn uniformly from [0, clip duration], wrapping around the clip boundary. This procedure preserves within-species call rates and inter-call timing while eliminating any cross-species correlation in call timing. A one-sided permutation *p*-value was computed as the fraction of permuted mean reply probabilities that equaled or exceeded the observed value. All 12 ordered off-diagonal species pairs were tested simultaneously, and p-values were Bonferroni-corrected with *α* = 0.05.

### 1.10 Generative AI Usage

Generative AI (Claude Sonnet 4.6) was used to produce initial drafts of some text in the Online Methods section. This text was revised for correctness and clarity by the authors.

