## Supplemental Materials for "BirdCODE: Detecting bird communication at scale"

### Supplementary Materials

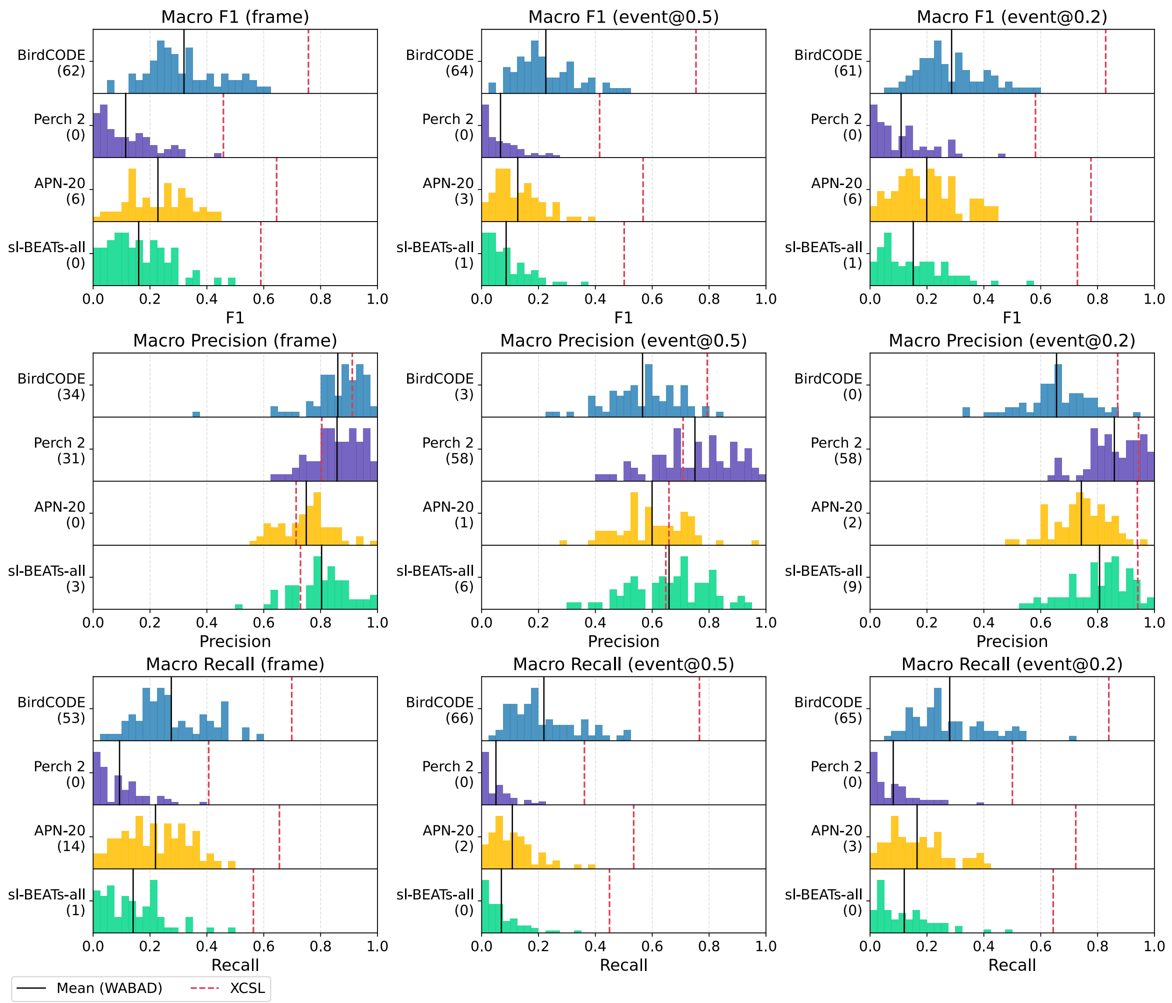


**Figure S1.** Performance comparison using supplementary metrics for BirdCODE against baseline methods using sliding-window classifiers. The number in parentheses below each model name is the number of WABAD datasets where the model achieved the top score in that metric. Histogram of performance on WABAD datasets, with lines for WABAD mean and XCSL.

Table S1. Per-dataset mAP (frame): Results for each of the 68 WABAD datasets, average across WABAD, and results for XCSL. The best value in each column is shown in bold.

| Model | ARD | BAM | BIAL | BMT | BOLIN | BRCAS | BRE | BUR | CARI | CAT | CB | CLH | COU | CRUZ | DEVA |
|---|---|---|---|---|---|---|---|---|---|---|---|---|---|---|---|
| sl-BEATs-all | 0.19 | 0.10 | 0.31 | 0.11 | 0.10 | 0.13 | 0.18 | 0.12 | 0.11 | 0.29 | 0.15 | 0.41 | 0.27 | 0.12 | 0.13 |
| APN-20 | 0.40 | **0.36** | 0.49 | 0.34 | 0.21 | 0.38 | 0.36 | 0.38 | 0.30 | 0.43 | 0.28 | 0.49 | 0.50 | 0.31 | 0.41 |
| Perch 2 | 0.23 | 0.10 | 0.44 | 0.15 | 0.15 | 0.19 | 0.23 | 0.13 | 0.16 | 0.41 | 0.20 | 0.49 | 0.32 | 0.15 | 0.17 |
| BirdCODE | **0.53** | 0.25 | **0.60** | **0.42** | **0.24** | **0.60** | **0.45** | **0.44** | **0.44** | **0.58** | **0.40** | **0.59** | **0.64** | **0.35** | **0.53** |

| Model | DONG | DUNAS | DYOM | EMP | EVROS | FEU | FNCA | GLEN | GTLU | HAG | HAK | HAR | HONDO | HUAP | JUNCA |
|---|---|---|---|---|---|---|---|---|---|---|---|---|---|---|---|
| sl-BEATs-all | 0.10 | 0.29 | 0.13 | 0.41 | 0.27 | 0.19 | 0.11 | 0.30 | 0.09 | 0.21 | 0.11 | 0.24 | 0.24 | 0.24 | 0.24 |
| APN-20 | 0.30 | 0.43 | 0.40 | 0.57 | 0.38 | 0.39 | 0.30 | 0.49 | 0.32 | 0.43 | 0.21 | 0.43 | 0.39 | 0.43 | 0.42 |
| Perch 2 | 0.11 | 0.33 | 0.16 | 0.58 | 0.35 | 0.27 | 0.14 | 0.40 | 0.12 | 0.32 | 0.14 | 0.33 | 0.28 | 0.34 | 0.36 |
| BirdCODE | **0.40** | **0.73** | **0.42** | **0.68** | **0.41** | **0.58** | **0.41** | **0.57** | **0.49** | **0.57** | **0.36** | **0.58** | **0.40** | **0.56** | **0.51** |

| Model | KAR | KIB | LIM | MABI | MAPIMI | MARTI | MILLAN | MONTEB | MOPU | NAV | NL | OESF | OIO | OLIV | PETI |
|---|---|---|---|---|---|---|---|---|---|---|---|---|---|---|---|
| sl-BEATs-all | 0.34 | 0.10 | 0.11 | 0.33 | 0.28 | 0.15 | 0.26 | 0.22 | 0.19 | 0.26 | 0.20 | 0.44 | 0.18 | 0.40 | 0.23 |
| APN-20 | 0.50 | 0.29 | 0.28 | 0.41 | 0.42 | **0.23** | 0.27 | **0.45** | 0.42 | 0.36 | 0.38 | 0.56 | 0.47 | 0.45 | 0.44 |
| Perch 2 | 0.47 | 0.10 | 0.16 | 0.38 | 0.33 | 0.18 | 0.22 | 0.31 | 0.28 | 0.30 | 0.29 | 0.53 | 0.23 | 0.42 | 0.30 |
| BirdCODE | **0.64** | **0.30** | **0.29** | **0.49** | **0.51** | 0.22 | **0.34** | 0.41 | **0.48** | **0.46** | **0.49** | **0.77** | **0.61** | **0.66** | **0.60** |

| Model | PGF | PINA | PITI | POZO | PUUL | QR | RBA | RFP | RGU | RME | ROTOK | SAL | SBN | SCHF | SCHG |
|---|---|---|---|---|---|---|---|---|---|---|---|---|---|---|---|
| sl-BEATs-all | 0.07 | 0.21 | 0.13 | 0.15 | 0.15 | 0.33 | 0.30 | 0.28 | 0.35 | 0.39 | 0.04 | 0.18 | 0.17 | 0.51 | 0.19 |
| APN-20 | **0.32** | 0.40 | 0.44 | 0.30 | 0.39 | 0.50 | 0.47 | 0.51 | 0.53 | 0.58 | **0.15** | 0.32 | 0.30 | 0.55 | 0.38 |
| Perch 2 | 0.08 | 0.32 | 0.29 | 0.23 | 0.22 | 0.43 | 0.42 | 0.38 | 0.45 | 0.54 | 0.09 | 0.20 | 0.16 | 0.56 | 0.27 |
| BirdCODE | 0.21 | **0.54** | **0.51** | **0.36** | **0.58** | **0.66** | **0.67** | **0.73** | **0.60** | **0.72** | 0.14 | **0.38** | **0.35** | **0.61** | **0.45** |

| Model | SD | SITH | SLOB | SPMCO | TAM | UNI | VER | VIL | Avg | XCSL |
|---|---|---|---|---|---|---|---|---|---|---|
| sl-BEATs-all | 0.19 | 0.22 | 0.25 | 0.13 | 0.17 | 0.17 | 0.28 | 0.33 | 0.22 | 0.66 |
| APN-20 | 0.33 | 0.34 | 0.39 | 0.33 | 0.42 | 0.26 | 0.43 | 0.49 | 0.39 | 0.68 |
| Perch 2 | 0.21 | 0.28 | 0.35 | 0.20 | 0.31 | 0.21 | 0.30 | 0.39 | 0.28 | 0.65 |
| BirdCODE | **0.46** | **0.41** | **0.61** | **0.48** | **0.58** | **0.34** | **0.49** | **0.60** | **0.49** | **0.84** |

Table S2. Per-dataset mAP (event@0.5): Results for each of the 68 WABAD datasets, average across WABAD, and results for XCSL. The best value in each column is shown in bold.

| Model | ARD | BAM | BIAL | BMT | BOLIN | BRCAS | BRE | BUR | CARI | CAT | CB | CLH | COU | CRUZ | DEVA |
|---|---|---|---|---|---|---|---|---|---|---|---|---|---|---|---|
| sl-BEATs-all | 0.09 | 0.04 | 0.17 | 0.04 | 0.05 | 0.04 | 0.09 | 0.04 | 0.07 | 0.10 | 0.04 | 0.19 | 0.12 | 0.07 | 0.03 |
| APN-20 | 0.18 | **0.16** | **0.33** | 0.10 | 0.06 | 0.09 | 0.18 | 0.13 | 0.12 | 0.11 | 0.09 | 0.22 | 0.26 | 0.16 | 0.13 |
| Perch 2 | 0.11 | 0.05 | 0.30 | 0.06 | 0.04 | 0.07 | 0.09 | 0.04 | 0.07 | 0.11 | 0.07 | **0.26** | 0.17 | 0.08 | 0.09 |
| BirdCODE | **0.33** | 0.13 | 0.32 | **0.23** | **0.15** | **0.32** | **0.26** | **0.19** | **0.24** | **0.32** | **0.23** | 0.22 | **0.47** | **0.24** | **0.37** |

| Model | DONG | DUNAS | DYOM | EMP | EVROS | FEU | FNCA | GLEN | GTLU | HAG | HAK | HAR | HONDO | HUAP | JUNCA |
|---|---|---|---|---|---|---|---|---|---|---|---|---|---|---|---|
| sl-BEATs-all | 0.02 | 0.10 | 0.05 | 0.34 | 0.15 | 0.07 | 0.05 | 0.18 | 0.04 | 0.08 | 0.01 | 0.11 | 0.09 | 0.07 | 0.08 |
| APN-20 | 0.08 | 0.20 | 0.10 | 0.44 | 0.23 | 0.14 | 0.19 | **0.34** | 0.13 | 0.17 | 0.02 | 0.23 | 0.12 | 0.17 | 0.14 |
| Perch 2 | 0.02 | 0.17 | 0.06 | 0.50 | 0.21 | 0.12 | 0.07 | 0.25 | 0.03 | 0.12 | 0.03 | 0.17 | 0.08 | 0.14 | 0.07 |
| BirdCODE | **0.16** | **0.54** | **0.23** | **0.56** | **0.24** | **0.31** | **0.30** | 0.31 | **0.29** | **0.24** | **0.10** | **0.39** | **0.24** | **0.37** | **0.36** |

| Model | KAR | KIB | LIM | MABI | MAPIMI | MARTI | MILLAN | MONTEB | MOPU | NAV | NL | OESF | OIO | OLIV | PETI |
|---|---|---|---|---|---|---|---|---|---|---|---|---|---|---|---|
| sl-BEATs-all | 0.21 | 0.04 | 0.02 | 0.24 | 0.15 | 0.05 | 0.10 | 0.12 | 0.08 | 0.10 | 0.14 | 0.33 | 0.11 | 0.14 | 0.15 |
| APN-20 | **0.38** | **0.13** | 0.06 | 0.27 | 0.20 | 0.10 | 0.14 | 0.22 | 0.19 | 0.14 | 0.20 | 0.42 | 0.24 | 0.15 | 0.26 |
| Perch 2 | 0.35 | 0.04 | 0.02 | 0.26 | 0.16 | 0.08 | 0.10 | 0.16 | 0.10 | 0.16 | 0.21 | 0.38 | 0.13 | 0.14 | 0.20 |
| BirdCODE | 0.37 | **0.13** | **0.12** | **0.30** | **0.32** | **0.15** | **0.32** | **0.29** | **0.34** | **0.28** | **0.37** | **0.57** | **0.40** | **0.38** | **0.47** |

| Model | PGF | PINA | PITI | POZO | PUUL | QR | RBA | RFP | RGU | RME | ROTOK | SAL | SBN | SCHF | SCHG |
|---|---|---|---|---|---|---|---|---|---|---|---|---|---|---|---|
| sl-BEATs-all | 0.02 | 0.06 | 0.05 | 0.03 | 0.03 | 0.12 | 0.19 | 0.15 | 0.23 | 0.24 | 0.02 | 0.08 | 0.03 | 0.26 | 0.06 |
| APN-20 | **0.16** | 0.12 | 0.18 | 0.06 | 0.09 | 0.23 | 0.28 | 0.28 | 0.41 | 0.37 | 0.04 | 0.12 | 0.05 | 0.30 | 0.10 |
| Perch 2 | 0.02 | 0.10 | 0.12 | 0.06 | 0.07 | 0.18 | 0.25 | 0.21 | 0.33 | 0.39 | 0.02 | 0.07 | 0.02 | 0.23 | 0.08 |
| BirdCODE | 0.13 | **0.24** | **0.32** | **0.29** | **0.23** | **0.49** | **0.54** | **0.53** | **0.58** | **0.60** | **0.07** | **0.24** | **0.24** | **0.34** | **0.15** |

| Model | SD | SITH | SLOB | SPMCO | TAM | UNI | VER | VIL | Avg | XCSL |
|---|---|---|---|---|---|---|---|---|---|---|
| sl-BEATs-all | 0.06 | 0.11 | 0.08 | 0.04 | 0.08 | 0.07 | 0.06 | 0.11 | 0.10 | 0.41 |
| APN-20 | 0.11 | 0.13 | 0.16 | 0.13 | 0.22 | 0.12 | 0.10 | 0.18 | 0.18 | 0.47 |
| Perch 2 | 0.07 | 0.16 | 0.15 | 0.10 | 0.18 | 0.11 | 0.07 | 0.15 | 0.14 | 0.42 |
| BirdCODE | **0.32** | **0.25** | **0.36** | **0.25** | **0.43** | **0.25** | **0.22** | **0.31** | **0.31** | **0.71** |

**Table S3.** Per-dataset mAP (event@0.2): Results for each of the 68 WABAD datasets, average across WABAD, and results for XCSL. The best value in each column is shown in bold.

| Model | ARD | BAM | BIAL | BMT | BOLIN | BRCAS | BRE | BUR | CARI | CAT | CB | CLH | COU | CRUZ | DEVA |
|---|---|---|---|---|---|---|---|---|---|---|---|---|---|---|---|
| sl-BEATs-all | 0.16 | 0.08 | 0.26 | 0.06 | 0.09 | 0.09 | 0.14 | 0.08 | 0.09 | 0.23 | 0.15 | 0.32 | 0.27 | 0.11 | 0.13 |
| APN-20 | 0.23 | **0.21** | **0.39** | 0.14 | **0.15** | 0.20 | 0.22 | 0.18 | 0.19 | 0.28 | 0.21 | 0.34 | 0.37 | 0.22 | 0.22 |
| Perch 2 | 0.15 | 0.08 | **0.39** | 0.08 | 0.13 | 0.14 | 0.18 | 0.08 | 0.13 | 0.27 | 0.19 | **0.37** | 0.31 | 0.12 | 0.13 |
| BirdCODE | **0.32** | 0.16 | 0.34 | **0.24** | **0.15** | **0.32** | **0.34** | **0.21** | **0.29** | **0.45** | **0.28** | 0.32 | **0.53** | **0.28** | **0.32** |

| Model | DONG | DUNAS | DYOM | EMP | EVROS | FEU | FNCA | GLEN | GTLU | HAG | HAK | HAR | HONDO | HUAP | JUNCA |
|---|---|---|---|---|---|---|---|---|---|---|---|---|---|---|---|
| sl-BEATs-all | 0.05 | 0.27 | 0.09 | 0.45 | 0.26 | 0.09 | 0.09 | 0.31 | 0.06 | 0.12 | 0.06 | 0.21 | 0.17 | 0.18 | 0.21 |
| APN-20 | 0.09 | 0.43 | 0.20 | 0.57 | **0.38** | 0.16 | 0.27 | **0.43** | 0.21 | 0.22 | 0.05 | 0.28 | 0.24 | 0.34 | 0.27 |
| Perch 2 | 0.04 | 0.37 | 0.10 | 0.59 | 0.31 | 0.16 | 0.12 | 0.36 | 0.05 | 0.19 | 0.04 | 0.23 | 0.21 | 0.28 | 0.24 |
| BirdCODE | **0.16** | **0.59** | **0.28** | **0.69** | 0.26 | **0.38** | **0.38** | 0.36 | **0.26** | **0.29** | **0.15** | **0.41** | **0.27** | **0.42** | **0.44** |

| Model | KAR | KIB | LIM | MABI | MAPIMI | MARTI | MILLAN | MONTEB | MOPU | NAV | NL | OESF | OIO | OLIV | PETI |
|---|---|---|---|---|---|---|---|---|---|---|---|---|---|---|---|
| sl-BEATs-all | 0.33 | 0.08 | 0.03 | 0.35 | 0.23 | 0.16 | 0.21 | 0.21 | 0.16 | 0.21 | 0.18 | 0.42 | 0.17 | 0.31 | 0.30 |
| APN-20 | 0.47 | **0.17** | 0.15 | **0.42** | 0.36 | 0.18 | 0.26 | 0.32 | 0.26 | 0.24 | 0.31 | 0.54 | 0.33 | 0.32 | 0.48 |
| Perch 2 | **0.53** | 0.07 | 0.12 | 0.39 | 0.31 | 0.20 | 0.17 | 0.31 | 0.20 | 0.25 | 0.27 | 0.48 | 0.25 | 0.37 | 0.36 |
| BirdCODE | 0.44 | 0.15 | **0.18** | 0.35 | **0.51** | **0.21** | **0.37** | **0.37** | **0.40** | **0.31** | **0.46** | **0.61** | **0.49** | **0.49** | **0.49** |

| Model | PGF | PINA | PITI | POZO | PUUL | QR | RBA | RFP | RGU | RME | ROTOK | SAL | SBN | SCHF | SCHG |
|---|---|---|---|---|---|---|---|---|---|---|---|---|---|---|---|
| sl-BEATs-all | 0.03 | 0.13 | 0.08 | 0.08 | 0.08 | 0.31 | 0.23 | 0.29 | 0.36 | 0.40 | 0.03 | 0.17 | 0.16 | **0.55** | 0.12 |
| APN-20 | **0.18** | 0.25 | 0.32 | 0.19 | 0.16 | 0.46 | 0.37 | 0.45 | 0.54 | 0.51 | **0.14** | 0.23 | 0.25 | 0.45 | 0.20 |
| Perch 2 | 0.04 | 0.19 | 0.22 | 0.13 | 0.14 | 0.38 | 0.35 | 0.41 | 0.47 | 0.56 | 0.11 | 0.17 | 0.20 | **0.55** | 0.16 |
| BirdCODE | 0.17 | **0.28** | **0.38** | **0.33** | **0.28** | **0.58** | **0.56** | **0.60** | **0.67** | **0.68** | 0.12 | **0.28** | **0.32** | 0.48 | **0.22** |

| Model | SD | SITH | SLOB | SPMCO | TAM | UNI | VER | VIL | Avg | XCSL |
|---|---|---|---|---|---|---|---|---|---|---|
| sl-BEATs-all | 0.16 | 0.20 | 0.17 | 0.10 | 0.20 | 0.18 | 0.17 | 0.23 | 0.19 | 0.70 |
| APN-20 | 0.24 | **0.28** | 0.25 | 0.21 | 0.37 | 0.22 | 0.21 | 0.31 | 0.28 | 0.75 |
| Perch 2 | 0.14 | **0.28** | 0.23 | 0.18 | 0.26 | 0.22 | 0.17 | 0.28 | 0.24 | 0.70 |
| BirdCODE | **0.30** | 0.25 | **0.37** | **0.29** | **0.42** | **0.26** | **0.28** | **0.41** | **0.36** | **0.80** |

**Table S4.** Per-dataset Macro F1 (frame): Results for each of the 68 WABAD datasets, average across WABAD, and results for XCSL. The best value in each column is shown in bold.

| Model | ARD | BAM | BIAL | BMT | BOLIN | BRCAS | BRE | BUR | CARI | CAT | CB | CLH | COU | CRUZ | DEVA |
|---|---|---|---|---|---|---|---|---|---|---|---|---|---|---|---|
| sl-BEATs-all | 0.10 | 0.02 | 0.28 | 0.09 | 0.10 | 0.09 | 0.08 | 0.04 | 0.06 | 0.29 | 0.14 | 0.37 | 0.22 | 0.05 | 0.10 |
| APN-20 | 0.18 | 0.05 | 0.41 | 0.15 | 0.14 | 0.18 | 0.14 | 0.09 | 0.15 | 0.34 | 0.18 | 0.39 | 0.22 | 0.16 | 0.13 |
| Perch 2 | 0.02 | 0.02 | 0.28 | 0.04 | 0.06 | 0.04 | 0.02 | 0.01 | 0.04 | 0.28 | 0.09 | 0.30 | 0.18 | 0.02 | 0.04 |
| BirdCODE | **0.30** | **0.06** | **0.56** | **0.23** | **0.21** | **0.33** | **0.28** | **0.23** | **0.26** | **0.42** | **0.24** | **0.54** | **0.28** | **0.27** | **0.33** |

| Model | DONG | DUNAS | DYOM | EMP | EVROS | FEU | FNCA | GLEN | GTLU | HAG | HAK | HAR | HONDO | HUAP | JUNCA |
|---|---|---|---|---|---|---|---|---|---|---|---|---|---|---|---|
| sl-BEATs-all | 0.04 | 0.17 | 0.07 | 0.21 | 0.24 | 0.13 | 0.06 | 0.28 | 0.03 | 0.19 | 0.00 | 0.19 | 0.25 | 0.09 | 0.26 |
| APN-20 | 0.10 | 0.30 | 0.12 | 0.38 | **0.30** | 0.14 | 0.13 | 0.39 | 0.10 | 0.31 | 0.06 | **0.26** | 0.28 | 0.23 | 0.27 |
| Perch 2 | 0.01 | 0.12 | 0.05 | 0.16 | 0.18 | 0.10 | 0.02 | 0.27 | 0.01 | 0.17 | 0.00 | 0.16 | 0.16 | 0.05 | 0.13 |
| BirdCODE | **0.17** | **0.53** | **0.20** | **0.41** | 0.27 | **0.27** | **0.15** | **0.48** | **0.15** | **0.34** | **0.28** | 0.25 | **0.30** | **0.34** | **0.37** |

| Model | KAR | KIB | LIM | MABI | MAPIMI | MARTI | MILLAN | MONTEB | MOPU | NAV | NL | OESF | OIO | OLIV | PETI |
|---|---|---|---|---|---|---|---|---|---|---|---|---|---|---|---|
| sl-BEATs-all | 0.29 | 0.03 | 0.01 | 0.33 | 0.20 | 0.10 | 0.11 | 0.14 | 0.16 | 0.30 | 0.17 | 0.44 | 0.06 | 0.36 | 0.13 |
| APN-20 | 0.34 | 0.09 | **0.13** | 0.27 | 0.27 | 0.14 | 0.21 | **0.29** | 0.23 | 0.31 | 0.27 | 0.44 | 0.14 | 0.30 | **0.26** |
| Perch 2 | 0.21 | 0.00 | 0.03 | 0.25 | 0.09 | 0.12 | 0.08 | 0.20 | 0.12 | 0.24 | 0.20 | 0.31 | 0.03 | 0.28 | 0.05 |
| BirdCODE | **0.54** | **0.18** | **0.13** | **0.35** | **0.37** | **0.22** | **0.25** | 0.27 | **0.28** | **0.42** | **0.32** | **0.58** | **0.30** | **0.47** | 0.25 |

| Model | PGF | PINA | PITI | POZO | PUUL | QR | RBA | RFP | RGU | RME | ROTOK | SAL | SBN | SCHF | SCHG |
|---|---|---|---|---|---|---|---|---|---|---|---|---|---|---|---|
| sl-BEATs-all | 0.01 | 0.24 | 0.01 | 0.10 | 0.07 | 0.21 | 0.16 | 0.18 | 0.24 | 0.25 | 0.04 | 0.13 | 0.09 | 0.50 | 0.17 |
| APN-20 | 0.02 | 0.35 | **0.36** | 0.12 | 0.15 | 0.25 | 0.28 | 0.33 | 0.34 | 0.45 | 0.03 | 0.16 | 0.05 | 0.42 | **0.25** |
| Perch 2 | 0.00 | 0.19 | 0.18 | 0.05 | 0.07 | 0.12 | 0.13 | 0.11 | 0.09 | 0.17 | 0.03 | 0.03 | 0.03 | 0.43 | 0.16 |
| BirdCODE | **0.05** | **0.51** | 0.25 | **0.20** | **0.25** | **0.47** | **0.46** | **0.50** | **0.48** | **0.55** | **0.14** | **0.30** | **0.21** | **0.62** | 0.19 |

| Model | SD | SITH | SLOB | SPMCO | TAM | UNI | VER | VIL | Avg | XCSL |
|---|---|---|---|---|---|---|---|---|---|---|
| sl-BEATs-all | 0.10 | 0.21 | 0.27 | 0.10 | 0.17 | 0.12 | 0.22 | 0.23 | 0.16 | 0.59 |
| APN-20 | 0.18 | 0.20 | 0.33 | 0.19 | 0.32 | 0.16 | 0.26 | 0.31 | 0.23 | 0.65 |
| Perch 2 | 0.06 | 0.14 | 0.22 | 0.03 | 0.06 | 0.06 | 0.05 | 0.13 | 0.11 | 0.46 |
| BirdCODE | **0.24** | **0.33** | **0.40** | **0.20** | **0.43** | **0.25** | **0.38** | **0.34** | **0.32** | **0.76** |

**Table S5.** Per-dataset Macro F1 (event@0.5): Results for each of the 68 WABAD datasets, average across WABAD, and results for XCSL. The best value in each column is shown in bold.

| Model | ARD | BAM | BIAL | BMT | BOLIN | BRCAS | BRE | BUR | CARI | CAT | CB | CLH | COU | CRUZ | DEVA |
|---|---|---|---|---|---|---|---|---|---|---|---|---|---|---|---|
| sl-BEATs-all | 0.06 | 0.02 | 0.19 | 0.04 | 0.03 | 0.05 | 0.03 | 0.01 | 0.05 | 0.15 | 0.07 | 0.21 | 0.15 | 0.03 | 0.03 |
| APN-20 | 0.10 | **0.04** | 0.32 | 0.07 | 0.06 | 0.07 | 0.07 | 0.04 | 0.08 | 0.17 | 0.08 | 0.22 | 0.15 | 0.10 | 0.07 |
| Perch 2 | 0.02 | 0.02 | 0.24 | 0.02 | 0.02 | 0.03 | 0.01 | 0.00 | 0.02 | 0.13 | 0.03 | 0.19 | 0.12 | 0.02 | 0.01 |
| BirdCODE | **0.21** | **0.04** | **0.39** | **0.16** | **0.14** | **0.22** | **0.15** | **0.13** | **0.20** | **0.31** | **0.20** | **0.29** | **0.24** | **0.22** | **0.28** |

| Model | DONG | DUNAS | DYOM | EMP | EVROS | FEU | FNCA | GLEN | GTLU | HAG | HAK | HAR | HONDO | HUAP | JUNCA |
|---|---|---|---|---|---|---|---|---|---|---|---|---|---|---|---|
| sl-BEATs-all | 0.01 | 0.09 | 0.05 | 0.19 | 0.15 | 0.06 | 0.02 | 0.19 | 0.01 | 0.08 | 0.00 | 0.13 | 0.11 | 0.03 | 0.07 |
| APN-20 | 0.03 | 0.18 | 0.08 | 0.26 | **0.18** | 0.07 | 0.08 | 0.25 | 0.06 | 0.15 | 0.02 | **0.17** | 0.08 | 0.12 | 0.07 |
| Perch 2 | 0.00 | 0.07 | 0.03 | 0.14 | 0.12 | 0.06 | 0.00 | 0.19 | 0.01 | 0.07 | 0.00 | 0.11 | 0.04 | 0.02 | 0.04 |
| BirdCODE | **0.09** | **0.45** | **0.09** | **0.31** | 0.16 | **0.17** | **0.12** | **0.32** | **0.10** | **0.19** | **0.09** | 0.13 | **0.18** | **0.27** | **0.27** |

| Model | KAR | KIB | LIM | MABI | MAPIMI | MARTI | MILLAN | MONTEB | MOPU | NAV | NL | OESF | OIO | OLIV | PETI |
|---|---|---|---|---|---|---|---|---|---|---|---|---|---|---|---|
| sl-BEATs-all | 0.20 | 0.02 | 0.00 | **0.27** | 0.04 | 0.03 | 0.04 | 0.07 | 0.08 | 0.16 | 0.13 | 0.35 | 0.03 | 0.21 | 0.09 |
| APN-20 | 0.25 | 0.05 | 0.04 | 0.20 | 0.15 | 0.07 | **0.14** | 0.16 | 0.13 | 0.16 | 0.24 | 0.40 | 0.10 | 0.10 | **0.21** |
| Perch 2 | 0.14 | 0.00 | 0.00 | 0.21 | 0.02 | 0.06 | 0.03 | 0.12 | 0.09 | 0.15 | 0.15 | 0.23 | 0.01 | 0.08 | 0.04 |
| BirdCODE | **0.31** | **0.09** | **0.10** | 0.23 | **0.31** | **0.14** | **0.14** | **0.18** | **0.15** | **0.29** | **0.25** | **0.52** | **0.21** | **0.32** | **0.21** |

| Model | PGF | PINA | PITI | POZO | PUUL | QR | RBA | RFP | RGU | RME | ROTOK | SAL | SBN | SCHF | SCHG |
|---|---|---|---|---|---|---|---|---|---|---|---|---|---|---|---|
| sl-BEATs-all | 0.00 | 0.09 | 0.00 | 0.01 | 0.02 | 0.07 | 0.09 | 0.11 | 0.11 | 0.13 | 0.02 | 0.05 | 0.02 | 0.29 | 0.09 |
| APN-20 | 0.00 | 0.15 | 0.16 | 0.03 | 0.06 | 0.10 | 0.13 | 0.20 | 0.33 | 0.18 | 0.01 | 0.07 | 0.01 | 0.26 | **0.12** |
| Perch 2 | 0.00 | 0.09 | 0.09 | 0.02 | 0.03 | 0.05 | 0.06 | 0.06 | 0.10 | 0.10 | 0.01 | 0.01 | 0.00 | 0.27 | 0.09 |
| BirdCODE | **0.04** | **0.34** | **0.18** | **0.13** | **0.16** | **0.36** | **0.36** | **0.48** | **0.46** | **0.39** | **0.08** | **0.18** | **0.20** | **0.44** | **0.12** |

| Model | SD | SITH | SLOB | SPMCO | TAM | UNI | VER | VIL | Avg | XCSL |
|---|---|---|---|---|---|---|---|---|---|---|
| sl-BEATs-all | 0.06 | 0.17 | 0.14 | 0.03 | 0.09 | 0.09 | 0.07 | 0.09 | 0.09 | 0.50 |
| APN-20 | 0.09 | 0.13 | 0.19 | 0.09 | 0.19 | 0.12 | 0.08 | 0.14 | 0.13 | 0.57 |
| Perch 2 | 0.04 | 0.08 | 0.12 | 0.01 | 0.02 | 0.06 | 0.02 | 0.06 | 0.07 | 0.41 |
| BirdCODE | **0.20** | **0.21** | **0.30** | **0.13** | **0.39** | **0.20** | **0.19** | **0.25** | **0.23** | **0.75** |

**Table S6.** Per-dataset Macro F1 (event@0.2): Results for each of the 68 WABAD datasets, average across WABAD, and results for XCSL. The best value in each column is shown in bold.

| Model | ARD | BAM | BIAL | BMT | BOLIN | BRCAS | BRE | BUR | CARI | CAT | CB | CLH | COU | CRUZ | DEVA |
|---|---|---|---|---|---|---|---|---|---|---|---|---|---|---|---|
| sl-BEATs-all | 0.07 | 0.03 | 0.29 | 0.06 | 0.06 | 0.07 | 0.07 | 0.04 | 0.06 | 0.27 | 0.14 | 0.35 | 0.26 | 0.05 | 0.10 |
| APN-20 | 0.13 | 0.05 | 0.36 | 0.08 | 0.12 | 0.12 | 0.09 | 0.08 | 0.12 | 0.30 | 0.15 | **0.39** | 0.26 | 0.13 | 0.13 |
| Perch 2 | 0.02 | 0.03 | 0.29 | 0.02 | 0.05 | 0.03 | 0.01 | 0.00 | 0.03 | 0.24 | 0.08 | 0.30 | 0.23 | 0.02 | 0.04 |
| BirdCODE | **0.27** | **0.08** | **0.37** | **0.17** | **0.19** | **0.32** | **0.23** | **0.17** | **0.26** | **0.46** | **0.24** | 0.38 | **0.32** | **0.25** | **0.27** |

| Model | DONG | DUNAS | DYOM | EMP | EVROS | FEU | FNCA | GLEN | GTLU | HAG | HAK | HAR | HONDO | HUAP | JUNCA |
|---|---|---|---|---|---|---|---|---|---|---|---|---|---|---|---|
| sl-BEATs-all | 0.03 | 0.20 | 0.07 | 0.25 | 0.24 | 0.09 | 0.05 | 0.30 | 0.01 | 0.13 | 0.00 | 0.18 | 0.17 | 0.06 | 0.22 |
| APN-20 | 0.04 | 0.28 | 0.14 | **0.41** | **0.29** | 0.06 | 0.12 | 0.37 | 0.07 | 0.17 | 0.03 | 0.21 | 0.19 | 0.21 | 0.25 |
| Perch 2 | 0.00 | 0.14 | 0.05 | 0.18 | 0.18 | 0.06 | 0.02 | 0.29 | 0.01 | 0.13 | 0.00 | 0.14 | 0.11 | 0.04 | 0.11 |
| BirdCODE | **0.11** | **0.51** | **0.17** | **0.41** | 0.23 | **0.24** | **0.16** | **0.39** | **0.13** | **0.24** | **0.19** | **0.22** | **0.21** | **0.33** | **0.31** |

| Model | KAR | KIB | LIM | MABI | MAPIMI | MARTI | MILLAN | MONTEB | MOPU | NAV | NL | OESF | OIO | OLIV | PETI |
|---|---|---|---|---|---|---|---|---|---|---|---|---|---|---|---|
| sl-BEATs-all | 0.32 | 0.03 | 0.01 | **0.37** | 0.21 | 0.10 | 0.08 | 0.10 | 0.11 | 0.25 | 0.14 | 0.45 | 0.05 | 0.34 | 0.16 |
| APN-20 | 0.39 | 0.09 | 0.11 | 0.27 | 0.25 | 0.13 | 0.20 | **0.22** | 0.17 | 0.23 | 0.30 | 0.44 | 0.15 | 0.25 | **0.30** |
| Perch 2 | 0.24 | 0.00 | 0.02 | 0.29 | 0.05 | 0.11 | 0.06 | 0.15 | 0.12 | 0.22 | 0.22 | 0.29 | 0.02 | 0.29 | 0.07 |
| BirdCODE | **0.44** | **0.13** | **0.14** | 0.35 | **0.43** | **0.21** | **0.23** | **0.22** | **0.22** | **0.34** | **0.33** | **0.56** | **0.32** | **0.41** | 0.28 |

| Model | PGF | PINA | PITI | POZO | PUUL | QR | RBA | RFP | RGU | RME | ROTOK | SAL | SBN | SCHF | SCHG |
|---|---|---|---|---|---|---|---|---|---|---|---|---|---|---|---|
| sl-BEATs-all | 0.01 | 0.19 | 0.01 | 0.06 | 0.05 | 0.23 | 0.15 | 0.20 | 0.32 | 0.26 | 0.04 | 0.11 | 0.08 | 0.55 | 0.16 |
| APN-20 | 0.00 | 0.28 | **0.26** | 0.05 | 0.11 | 0.23 | 0.21 | 0.36 | 0.38 | 0.42 | 0.02 | 0.16 | 0.06 | 0.44 | **0.20** |
| Perch 2 | 0.00 | 0.15 | 0.15 | 0.03 | 0.05 | 0.11 | 0.09 | 0.14 | 0.10 | 0.17 | 0.02 | 0.03 | 0.03 | 0.47 | 0.14 |
| BirdCODE | **0.07** | **0.34** | 0.23 | **0.20** | **0.19** | **0.44** | **0.40** | **0.55** | **0.48** | **0.50** | **0.11** | **0.23** | **0.24** | **0.58** | 0.19 |

| Model | SD | SITH | SLOB | SPMCO | TAM | UNI | VER | VIL | Avg | XCSL |
|---|---|---|---|---|---|---|---|---|---|---|
| sl-BEATs-all | 0.07 | 0.20 | 0.20 | 0.10 | 0.18 | 0.15 | 0.15 | 0.19 | 0.15 | 0.73 |
| APN-20 | 0.13 | 0.17 | 0.20 | 0.16 | 0.25 | 0.18 | 0.18 | 0.28 | 0.20 | 0.78 |
| Perch 2 | 0.04 | 0.16 | 0.18 | 0.02 | 0.06 | 0.07 | 0.04 | 0.13 | 0.11 | 0.58 |
| BirdCODE | **0.19** | **0.30** | **0.32** | **0.21** | **0.41** | **0.27** | **0.29** | **0.32** | **0.29** | **0.83** |

**Table S7**. Per-dataset Macro Precision (frame): Results for each of the 68 WABAD datasets, average across WABAD, and results for XCSL. The best value in each column is shown in bold.

| Model | ARD | BAM | BIAL | BMT | BOLIN | BRCAS | BRE | BUR | CARI | CAT | CB | CLH | COU | CRUZ | DEVA |
|---|---|---|---|---|---|---|---|---|---|---|---|---|---|---|---|
| sl-BEATs-all | 0.86 | **1.00** | 0.81 | 0.89 | 0.77 | 0.83 | 0.88 | 0.94 | 0.83 | 0.64 | 0.70 | 0.69 | 0.83 | 0.86 | 0.82 |
| APN-20 | 0.76 | 0.94 | 0.79 | 0.84 | 0.74 | 0.66 | 0.81 | 0.87 | 0.79 | 0.64 | 0.62 | 0.57 | 0.82 | 0.77 | 0.77 |
| Perch 2 | **0.94** | 0.96 | 0.84 | **0.96** | **0.81** | 0.92 | **0.96** | **0.97** | 0.83 | 0.70 | **0.75** | **0.77** | 0.89 | **0.94** | 0.90 |
| BirdCODE | 0.88 | 0.98 | **0.88** | 0.93 | **0.81** | **0.93** | 0.78 | 0.93 | **0.86** | **0.84** | 0.66 | 0.64 | **0.95** | 0.88 | **0.91** |

| Model | DONG | DUNAS | DYOM | EMP | EVROS | FEU | FNCA | GLEN | GTLU | HAG | HAK | HAR | HONDO | HUAP | JUNCA |
|---|---|---|---|---|---|---|---|---|---|---|---|---|---|---|---|
| sl-BEATs-all | 0.90 | 0.68 | 0.90 | 0.83 | 0.71 | 0.80 | 0.91 | 0.82 | **0.96** | 0.80 | 0.85 | 0.87 | 0.80 | 0.88 | 0.80 |
| APN-20 | 0.79 | 0.63 | 0.88 | 0.65 | 0.61 | 0.81 | 0.84 | 0.74 | 0.86 | 0.78 | 0.76 | 0.84 | 0.72 | 0.77 | 0.66 |
| Perch 2 | **0.97** | 0.81 | 0.84 | **0.84** | **0.86** | 0.87 | **0.96** | 0.78 | **0.96** | 0.84 | **1.00** | 0.90 | **0.85** | 0.93 | **0.90** |
| BirdCODE | 0.92 | **0.90** | **0.96** | 0.82 | 0.82 | **0.92** | 0.94 | **0.83** | 0.93 | **0.96** | 0.72 | **0.98** | **0.85** | **0.95** | 0.81 |

| Model | KAR | KIB | LIM | MABI | MAPIMI | MARTI | MILLAN | MONTEB | MOPU | NAV | NL | OESF | OIO | OLIV | PETI |
|---|---|---|---|---|---|---|---|---|---|---|---|---|---|---|---|
| sl-BEATs-all | 0.82 | 0.96 | 0.90 | 0.71 | 0.61 | 0.64 | 0.79 | 0.86 | 0.84 | 0.79 | 0.83 | 0.76 | 0.93 | 0.64 | 0.83 |
| APN-20 | 0.79 | 0.93 | 0.86 | 0.74 | 0.67 | 0.58 | 0.76 | 0.82 | 0.79 | 0.72 | 0.74 | 0.73 | 0.85 | 0.71 | 0.79 |
| Perch 2 | 0.84 | **0.99** | **0.94** | 0.79 | 0.66 | **0.74** | 0.80 | 0.88 | **0.92** | 0.80 | 0.73 | 0.82 | **0.96** | 0.72 | 0.89 |
| BirdCODE | **0.90** | 0.96 | 0.93 | **0.82** | **0.82** | 0.63 | **0.85** | **0.97** | 0.90 | **0.89** | **0.84** | **0.93** | 0.95 | **0.92** | **0.96** |

| Model | PGF | PINA | PITI | POZO | PUUL | QR | RBA | RFP | RGU | RME | ROTOK | SAL | SBN | SCHF | SCHG |
|---|---|---|---|---|---|---|---|---|---|---|---|---|---|---|---|
| sl-BEATs-all | 0.98 | 0.68 | **0.99** | 0.80 | 0.89 | 0.75 | 0.80 | 0.78 | 0.70 | 0.75 | 0.51 | 0.85 | 0.80 | 0.72 | 0.78 |
| APN-20 | **1.00** | 0.63 | 0.74 | 0.72 | 0.80 | 0.76 | 0.67 | 0.71 | 0.63 | 0.68 | 0.61 | 0.79 | 0.85 | 0.68 | 0.73 |
| Perch 2 | **1.00** | 0.75 | 0.87 | **0.86** | **0.92** | 0.88 | **0.81** | 0.80 | **0.91** | **0.86** | **0.64** | **0.95** | **0.92** | 0.77 | 0.75 |
| BirdCODE | 0.96 | **0.77** | 0.92 | 0.85 | 0.87 | **0.89** | 0.80 | **0.88** | 0.77 | 0.84 | 0.37 | 0.84 | 0.88 | **0.78** | **0.90** |

| Model | SD | SITH | SLOB | SPMCO | TAM | UNI | VER | VIL | Avg | XCSL |
|---|---|---|---|---|---|---|---|---|---|---|
| sl-BEATs-all | 0.86 | 0.65 | 0.69 | 0.83 | 0.71 | 0.76 | 0.81 | 0.78 | 0.80 | 0.73 |
| APN-20 | 0.78 | 0.62 | 0.61 | 0.76 | 0.58 | 0.79 | 0.80 | 0.77 | 0.75 | 0.71 |
| Perch 2 | 0.84 | **0.70** | 0.79 | 0.89 | 0.83 | **0.93** | **0.92** | 0.81 | **0.86** | 0.80 |
| BirdCODE | **0.89** | 0.69 | **0.86** | **0.95** | **0.84** | 0.81 | 0.76 | **0.93** | **0.86** | **0.91** |

**Table S8**. Per-dataset Macro Precision (event@0.5): Results for each of the 68 WABAD datasets, average across WABAD, and results for XCSL. The best value in each column is shown in bold.

| Model | ARD | BAM | BIAL | BMT | BOLIN | BRCAS | BRE | BUR | CARI | CAT | CB | CLH | COU | CRUZ | DEVA |
|---|---|---|---|---|---|---|---|---|---|---|---|---|---|---|---|
| sl-BEATs-all | 0.70 | 0.90 | 0.54 | 0.78 | 0.65 | 0.76 | 0.79 | 0.75 | 0.70 | 0.35 | 0.61 | 0.38 | 0.75 | 0.82 | 0.66 |
| APN-20 | 0.58 | 0.84 | 0.60 | 0.71 | 0.57 | 0.51 | 0.73 | 0.66 | 0.58 | **0.44** | 0.53 | 0.29 | 0.73 | 0.72 | 0.70 |
| Perch 2 | **0.90** | **0.92** | **0.70** | **0.91** | **0.68** | **0.86** | **0.94** | **0.90** | **0.75** | 0.43 | **0.64** | **0.47** | 0.75 | **0.94** | **0.81** |
| BirdCODE | 0.45 | 0.72 | 0.41 | 0.69 | 0.50 | 0.44 | 0.49 | 0.55 | 0.54 | 0.38 | 0.50 | 0.25 | **0.78** | 0.71 | 0.75 |

| Model | DONG | DUNAS | DYOM | EMP | EVROS | FEU | FNCA | GLEN | GTLU | HAG | HAK | HAR | HONDO | HUAP | JUNCA |
|---|---|---|---|---|---|---|---|---|---|---|---|---|---|---|---|
| sl-BEATs-all | 0.72 | 0.55 | **0.73** | 0.80 | 0.56 | 0.69 | 0.85 | 0.62 | 0.89 | 0.46 | 0.70 | 0.71 | 0.55 | 0.83 | 0.54 |
| APN-20 | 0.60 | 0.52 | 0.58 | 0.54 | 0.47 | 0.75 | 0.77 | 0.54 | 0.72 | 0.49 | 0.66 | 0.71 | 0.45 | 0.58 | 0.41 |
| Perch 2 | **0.92** | **0.74** | 0.68 | **0.82** | **0.67** | **0.80** | **0.94** | **0.64** | **0.93** | **0.52** | **0.93** | **0.80** | **0.60** | **0.85** | **0.66** |
| BirdCODE | 0.59 | 0.70 | 0.49 | 0.51 | 0.60 | 0.53 | 0.71 | 0.48 | 0.61 | 0.38 | 0.26 | 0.65 | 0.53 | 0.66 | 0.60 |

| Model | KAR | KIB | LIM | MABI | MAPIMI | MARTI | MILLAN | MONTEB | MOPU | NAV | NL | OESF | OIO | OLIV | PETI |
|---|---|---|---|---|---|---|---|---|---|---|---|---|---|---|---|
| sl-BEATs-all | 0.57 | 0.81 | 0.80 | 0.62 | 0.30 | 0.55 | 0.65 | **0.81** | 0.65 | 0.66 | **0.81** | 0.57 | 0.81 | 0.45 | 0.71 |
| APN-20 | 0.54 | 0.73 | 0.67 | 0.63 | 0.50 | 0.53 | 0.65 | 0.72 | 0.61 | 0.61 | 0.74 | 0.63 | 0.74 | 0.42 | 0.70 |
| Perch 2 | **0.60** | **0.96** | **0.90** | **0.68** | **0.52** | **0.69** | **0.76** | 0.77 | **0.81** | **0.71** | 0.68 | **0.64** | **0.88** | 0.40 | **0.85** |
| BirdCODE | 0.38 | 0.60 | 0.66 | 0.46 | 0.51 | 0.52 | 0.53 | 0.79 | 0.58 | 0.65 | 0.60 | 0.60 | 0.46 | **0.59** | 0.65 |

| Model | PGF | PINA | PITI | POZO | PUUL | QR | RBA | RFP | RGU | RME | ROTOK | SAL | SBN | SCHF | SCHG |
|---|---|---|---|---|---|---|---|---|---|---|---|---|---|---|---|
| sl-BEATs-all | 0.92 | 0.42 | **0.94** | 0.71 | 0.72 | 0.49 | 0.72 | 0.68 | 0.46 | 0.64 | 0.51 | 0.75 | 0.65 | 0.48 | **0.67** |
| APN-20 | 0.96 | 0.39 | 0.53 | 0.65 | 0.59 | 0.55 | 0.54 | 0.53 | 0.62 | 0.43 | **0.61** | 0.69 | 0.72 | 0.49 | 0.54 |
| Perch 2 | **1.00** | **0.53** | 0.68 | **0.83** | **0.76** | **0.71** | **0.76** | 0.68 | **0.96** | **0.82** | **0.61** | **0.90** | **0.82** | **0.52** | 0.63 |
| BirdCODE | 0.83 | 0.46 | 0.59 | 0.71 | 0.45 | 0.64 | 0.60 | **0.71** | 0.69 | 0.64 | 0.32 | 0.60 | 0.66 | 0.39 | 0.56 |

| Model | SD | SITH | SLOB | SPMCO | TAM | UNI | VER | VIL | Avg | XCSL |
|---|---|---|---|---|---|---|---|---|---|---|
| sl-BEATs-all | **0.80** | 0.51 | 0.51 | 0.63 | 0.60 | 0.68 | 0.69 | 0.51 | 0.66 | 0.65 |
| APN-20 | 0.72 | 0.46 | 0.45 | 0.60 | 0.42 | 0.75 | 0.63 | 0.53 | 0.60 | 0.66 |
| Perch 2 | **0.80** | **0.57** | **0.61** | **0.81** | **0.76** | **0.89** | **0.85** | **0.54** | **0.75** | 0.71 |
| BirdCODE | 0.73 | 0.41 | 0.52 | 0.61 | 0.64 | 0.59 | 0.54 | 0.53 | 0.57 | **0.79** |

**Table S9.** Per-dataset Macro Precision (event@0.2): Results for each of the 68 WABAD datasets, average across WABAD, and results for XCSL. The best value in each column is shown in bold.

| Model | ARD | BAM | BIAL | BMT | BOLIN | BRCAS | BRE | BUR | CARI | CAT | CB | CLH | COU | CRUZ | DEVA |
|---|---|---|---|---|---|---|---|---|---|---|---|---|---|---|---|
| sl-BEATs-all | 0.77 | **0.93** | 0.72 | 0.84 | 0.72 | 0.86 | **0.94** | 0.86 | 0.82 | 0.58 | **0.80** | 0.57 | 0.90 | 0.92 | 0.86 |
| APN-20 | 0.68 | 0.87 | 0.67 | 0.76 | 0.71 | 0.62 | 0.82 | 0.77 | 0.70 | 0.62 | 0.68 | 0.51 | 0.86 | 0.80 | 0.82 |
| Perch 2 | **0.91** | **0.93** | **0.80** | **0.90** | **0.78** | **0.87** | **0.94** | **0.92** | **0.84** | **0.69** | **0.80** | **0.65** | **0.92** | **0.97** | **0.93** |
| BirdCODE | 0.54 | 0.79 | 0.41 | 0.74 | 0.59 | 0.64 | 0.61 | 0.64 | 0.63 | 0.55 | 0.61 | 0.33 | 0.86 | 0.79 | 0.75 |

| Model | DONG | DUNAS | DYOM | EMP | EVROS | FEU | FNCA | GLEN | GTLU | HAG | HAK | HAR | HONDO | HUAP | JUNCA |
|---|---|---|---|---|---|---|---|---|---|---|---|---|---|---|---|
| sl-BEATs-all | 0.79 | 0.85 | **0.79** | 0.95 | 0.70 | 0.75 | 0.92 | **0.80** | **0.94** | 0.59 | 0.79 | 0.84 | 0.66 | 0.89 | 0.71 |
| APN-20 | 0.66 | 0.75 | 0.75 | 0.80 | 0.61 | 0.76 | 0.90 | 0.70 | 0.81 | 0.56 | 0.73 | 0.79 | 0.61 | 0.81 | 0.62 |
| Perch 2 | **0.95** | **0.94** | **0.79** | **0.98** | **0.85** | **0.81** | **1.00** | 0.77 | 0.93 | **0.67** | **1.00** | **0.86** | **0.76** | **0.92** | **0.78** |
| BirdCODE | 0.64 | 0.82 | 0.62 | 0.62 | 0.66 | 0.66 | 0.79 | 0.53 | 0.65 | 0.45 | 0.49 | 0.74 | 0.57 | 0.76 | 0.66 |

| Model | KAR | KIB | LIM | MABI | MAPIMI | MARTI | MILLAN | MONTEB | MOPU | NAV | NL | OESF | OIO | OLIV | PETI |
|---|---|---|---|---|---|---|---|---|---|---|---|---|---|---|---|
| sl-BEATs-all | 0.75 | 0.85 | **0.92** | **0.80** | 0.66 | 0.73 | 0.70 | **0.92** | 0.72 | 0.84 | **0.86** | 0.79 | 0.92 | 0.68 | 0.89 |
| APN-20 | 0.77 | 0.80 | 0.81 | 0.74 | **0.74** | 0.66 | 0.72 | 0.87 | 0.73 | 0.76 | **0.86** | 0.77 | 0.88 | 0.74 | 0.83 |
| Perch 2 | **0.79** | **0.97** | 0.89 | **0.80** | 0.67 | **0.83** | **0.83** | **0.92** | **0.86** | **0.88** | 0.84 | **0.84** | **0.96** | **0.79** | **0.93** |
| BirdCODE | 0.51 | 0.64 | 0.71 | 0.59 | 0.65 | 0.60 | 0.69 | 0.85 | 0.65 | 0.71 | 0.68 | 0.65 | 0.61 | 0.68 | 0.74 |

| Model | PGF | PINA | PITI | POZO | PUUL | QR | RBA | RFP | RGU | RME | ROTOK | SAL | SBN | SCHF | SCHG |
|---|---|---|---|---|---|---|---|---|---|---|---|---|---|---|---|
| sl-BEATs-all | **1.00** | 0.61 | **0.98** | 0.83 | 0.85 | 0.80 | **0.84** | 0.93 | 0.83 | 0.91 | 0.54 | 0.93 | 0.78 | 0.80 | **0.83** |
| APN-20 | 0.96 | 0.61 | 0.64 | 0.72 | 0.81 | 0.79 | 0.71 | 0.83 | 0.74 | 0.74 | 0.65 | 0.89 | 0.84 | 0.71 | 0.74 |
| Perch 2 | **1.00** | **0.66** | 0.81 | **0.89** | **0.86** | **0.85** | **0.84** | **0.94** | **0.96** | **0.96** | **0.67** | **0.99** | **0.91** | **0.81** | 0.78 |
| BirdCODE | 0.94 | 0.45 | 0.67 | 0.84 | 0.59 | 0.76 | 0.67 | 0.79 | 0.73 | 0.81 | 0.34 | 0.66 | 0.71 | 0.52 | 0.67 |

| Model | SD | SITH | SLOB | SPMCO | TAM | UNI | VER | VIL | Avg | XCSL |
|---|---|---|---|---|---|---|---|---|---|---|
| sl-BEATs-all | **0.85** | 0.63 | 0.64 | 0.87 | 0.82 | 0.87 | 0.89 | 0.71 | 0.81 | **0.94** |
| APN-20 | 0.80 | 0.59 | 0.48 | 0.72 | 0.62 | 0.85 | 0.86 | 0.75 | 0.74 | **0.94** |
| Perch 2 | 0.82 | **0.78** | **0.75** | **0.92** | **0.83** | **0.97** | **0.97** | **0.79** | **0.86** | **0.94** |
| BirdCODE | 0.74 | 0.49 | 0.60 | 0.76 | 0.67 | 0.71 | 0.70 | 0.64 | 0.66 | 0.87 |

**Table S10.** Per-dataset Macro Recall (frame): Results for each of the 68 WABAD datasets, average across WABAD, and results for XCSL. The best value in each column is shown in bold.

| Model | ARD | BAM | BIAL | BMT | BOLIN | BRCAS | BRE | BUR | CARI | CAT | CB | CLH | COU | CRUZ | DEVA |
|---|---|---|---|---|---|---|---|---|---|---|---|---|---|---|---|
| sl-BEATs-all | 0.07 | 0.02 | 0.24 | 0.07 | 0.08 | 0.06 | 0.07 | 0.03 | 0.05 | 0.33 | 0.13 | 0.35 | 0.21 | 0.04 | 0.07 |
| APN-20 | 0.16 | **0.05** | 0.35 | 0.12 | 0.12 | 0.16 | 0.17 | 0.07 | 0.14 | 0.36 | 0.20 | 0.39 | 0.21 | 0.14 | 0.11 |
| Perch 2 | 0.01 | 0.02 | 0.23 | 0.03 | 0.05 | 0.03 | 0.02 | 0.00 | 0.03 | 0.27 | 0.09 | 0.28 | 0.16 | 0.02 | 0.02 |
| BirdCODE | **0.21** | **0.05** | **0.47** | **0.18** | **0.16** | **0.23** | **0.33** | **0.18** | **0.22** | **0.38** | **0.23** | **0.53** | **0.26** | **0.24** | **0.27** |

| Model | DONG | DUNAS | DYOM | EMP | EVROS | FEU | FNCA | GLEN | GTLU | HAG | HAK | HAR | HONDO | HUAP | JUNCA |
|---|---|---|---|---|---|---|---|---|---|---|---|---|---|---|---|
| sl-BEATs-all | 0.02 | 0.16 | 0.05 | 0.20 | 0.22 | 0.11 | 0.05 | 0.24 | 0.02 | 0.14 | 0.00 | 0.13 | 0.21 | 0.07 | 0.22 |
| APN-20 | 0.08 | 0.30 | 0.08 | **0.44** | **0.31** | 0.13 | 0.09 | 0.37 | 0.07 | **0.24** | 0.04 | **0.20** | **0.32** | 0.21 | 0.30 |
| Perch 2 | 0.01 | 0.09 | 0.03 | 0.13 | 0.15 | 0.08 | 0.01 | 0.24 | 0.00 | 0.11 | 0.00 | 0.12 | 0.14 | 0.03 | 0.10 |
| BirdCODE | **0.12** | **0.46** | **0.13** | 0.39 | 0.24 | **0.20** | **0.10** | **0.42** | **0.11** | 0.23 | **0.29** | 0.18 | 0.25 | **0.28** | **0.32** |

| Model | KAR | KIB | LIM | MABI | MAPIMI | MARTI | MILLAN | MONTEB | MOPU | NAV | NL | OESF | OIO | OLIV | PETI |
|---|---|---|---|---|---|---|---|---|---|---|---|---|---|---|---|
| sl-BEATs-all | 0.22 | 0.01 | 0.01 | **0.30** | 0.18 | 0.09 | 0.14 | 0.11 | 0.12 | 0.25 | 0.17 | 0.41 | 0.04 | 0.34 | 0.11 |
| APN-20 | 0.27 | 0.06 | **0.11** | 0.22 | 0.30 | 0.17 | **0.28** | **0.27** | 0.20 | 0.29 | **0.27** | 0.42 | 0.12 | 0.27 | **0.27** |
| Perch 2 | 0.14 | 0.00 | 0.02 | 0.19 | 0.06 | 0.11 | 0.09 | 0.17 | 0.09 | 0.19 | 0.18 | 0.26 | 0.02 | 0.23 | 0.05 |
| BirdCODE | **0.43** | **0.13** | 0.10 | 0.26 | **0.32** | **0.19** | 0.24 | 0.23 | **0.22** | **0.36** | 0.25 | **0.47** | **0.22** | **0.38** | 0.18 |

| Model | PGF | PINA | PITI | POZO | PUUL | QR | RBA | RFP | RGU | RME | ROTOK | SAL | SBN | SCHF | SCHG |
|---|---|---|---|---|---|---|---|---|---|---|---|---|---|---|---|
| sl-BEATs-all | 0.00 | 0.19 | 0.01 | 0.10 | 0.04 | 0.21 | 0.18 | 0.17 | 0.22 | 0.24 | 0.04 | 0.11 | 0.10 | 0.49 | 0.13 |
| APN-20 | 0.01 | 0.34 | **0.36** | 0.16 | 0.10 | 0.25 | 0.31 | 0.34 | 0.36 | 0.48 | 0.02 | 0.14 | 0.03 | 0.42 | **0.21** |
| Perch 2 | 0.00 | 0.13 | 0.15 | 0.04 | 0.04 | 0.10 | 0.13 | 0.11 | 0.08 | 0.15 | 0.02 | 0.02 | 0.02 | 0.38 | 0.11 |
| BirdCODE | **0.04** | **0.45** | 0.20 | **0.20** | **0.17** | **0.41** | **0.46** | **0.43** | **0.47** | **0.54** | **0.22** | **0.26** | **0.16** | **0.58** | 0.14 |

| Model | SD | SITH | SLOB | SPMCO | TAM | UNI | VER | VIL | Avg | XCSL |
|---|---|---|---|---|---|---|---|---|---|---|
| sl-BEATs-all | 0.07 | 0.18 | 0.22 | 0.07 | 0.14 | 0.09 | 0.22 | 0.19 | 0.14 | 0.56 |
| APN-20 | **0.19** | 0.17 | 0.31 | **0.16** | 0.34 | 0.19 | 0.28 | **0.27** | 0.22 | 0.66 |
| Perch 2 | 0.04 | 0.10 | 0.17 | 0.02 | 0.04 | 0.05 | 0.04 | 0.09 | 0.09 | 0.41 |
| BirdCODE | **0.19** | **0.32** | **0.33** | 0.15 | **0.41** | **0.25** | **0.38** | 0.26 | **0.27** | **0.70** |

**Table S11.** Per-dataset Macro Recall (event@0.5): Results for each of the 68 WABAD datasets, average across WABAD, and results for XCSL. The best value in each column is shown in bold.

| Model | ARD | BAM | BIAL | BMT | BOLIN | BRCAS | BRE | BUR | CARI | CAT | CB | CLH | COU | CRUZ | DEVA |
|---|---|---|---|---|---|---|---|---|---|---|---|---|---|---|---|
| sl-BEATs-all | 0.04 | 0.02 | 0.14 | 0.03 | 0.02 | 0.03 | 0.02 | 0.01 | 0.03 | 0.12 | 0.05 | 0.20 | 0.11 | 0.02 | 0.02 |
| APN-20 | 0.07 | 0.03 | 0.27 | 0.05 | 0.04 | 0.05 | 0.08 | 0.03 | 0.06 | 0.13 | 0.07 | 0.21 | 0.12 | 0.08 | 0.05 |
| Perch 2 | 0.01 | 0.02 | 0.19 | 0.02 | 0.01 | 0.02 | 0.01 | 0.00 | 0.01 | 0.10 | 0.02 | 0.17 | 0.09 | 0.01 | 0.01 |
| BirdCODE | **0.19** | **0.05** | **0.41** | **0.14** | **0.12** | **0.18** | **0.14** | **0.12** | **0.19** | **0.31** | **0.18** | **0.40** | **0.22** | **0.19** | **0.25** |

| Model | DONG | DUNAS | DYOM | EMP | EVROS | FEU | FNCA | GLEN | GTLU | HAG | HAK | HAR | HONDO | HUAP | JUNCA |
|---|---|---|---|---|---|---|---|---|---|---|---|---|---|---|---|
| sl-BEATs-all | 0.01 | 0.07 | 0.04 | 0.18 | 0.14 | 0.04 | 0.01 | 0.16 | 0.00 | 0.06 | 0.00 | 0.10 | 0.09 | 0.02 | 0.05 |
| APN-20 | 0.02 | 0.15 | 0.06 | 0.26 | **0.16** | 0.06 | 0.05 | 0.23 | 0.04 | 0.12 | 0.01 | **0.13** | 0.09 | 0.10 | 0.05 |
| Perch 2 | 0.00 | 0.05 | 0.02 | 0.10 | 0.10 | 0.04 | 0.00 | 0.16 | 0.00 | 0.05 | 0.00 | 0.08 | 0.03 | 0.01 | 0.03 |
| BirdCODE | **0.08** | **0.41** | **0.08** | **0.29** | **0.16** | **0.14** | **0.10** | **0.36** | **0.09** | **0.16** | **0.07** | 0.11 | **0.19** | **0.26** | **0.24** |

| Model | KAR | KIB | LIM | MABI | MAPIMI | MARTI | MILLAN | MONTEB | MOPU | NAV | NL | OESF | OIO | OLIV | PETI |
|---|---|---|---|---|---|---|---|---|---|---|---|---|---|---|---|
| sl-BEATs-all | 0.16 | 0.01 | 0.00 | 0.23 | 0.03 | 0.02 | 0.03 | 0.04 | 0.07 | 0.12 | 0.10 | 0.33 | 0.02 | 0.17 | 0.08 |
| APN-20 | 0.22 | 0.04 | 0.03 | 0.15 | 0.14 | 0.06 | 0.13 | 0.13 | 0.12 | 0.12 | 0.21 | 0.38 | 0.08 | 0.08 | **0.18** |
| Perch 2 | 0.10 | 0.00 | 0.00 | 0.16 | 0.01 | 0.05 | 0.02 | 0.10 | 0.07 | 0.12 | 0.11 | 0.21 | 0.01 | 0.06 | 0.02 |
| BirdCODE | **0.37** | **0.10** | **0.09** | **0.24** | **0.30** | **0.13** | **0.15** | **0.16** | **0.17** | **0.26** | **0.24** | **0.51** | **0.19** | **0.32** | **0.18** |

| Model | PGF | PINA | PITI | POZO | PUUL | QR | RBA | RFP | RGU | RME | ROTOK | SAL | SBN | SCHF | SCHG |
|---|---|---|---|---|---|---|---|---|---|---|---|---|---|---|---|
| sl-BEATs-all | 0.00 | 0.07 | 0.00 | 0.01 | 0.01 | 0.05 | 0.07 | 0.09 | 0.11 | 0.10 | 0.01 | 0.04 | 0.01 | 0.26 | 0.06 |
| APN-20 | 0.00 | 0.13 | 0.15 | 0.02 | 0.04 | 0.08 | 0.11 | 0.17 | 0.33 | 0.15 | 0.01 | 0.05 | 0.00 | 0.22 | 0.09 |
| Perch 2 | 0.00 | 0.06 | 0.06 | 0.01 | 0.02 | 0.03 | 0.05 | 0.05 | 0.09 | 0.06 | 0.01 | 0.01 | 0.00 | 0.22 | 0.06 |
| BirdCODE | **0.04** | **0.37** | **0.17** | **0.11** | **0.12** | **0.32** | **0.34** | **0.45** | **0.48** | **0.37** | **0.09** | **0.16** | **0.20** | **0.52** | **0.10** |

| Model | SD | SITH | SLOB | SPMCO | TAM | UNI | VER | VIL | Avg | XCSL |
|---|---|---|---|---|---|---|---|---|---|---|
| sl-BEATs-all | 0.04 | 0.14 | 0.11 | 0.02 | 0.07 | 0.07 | 0.05 | 0.07 | 0.07 | 0.45 |
| APN-20 | 0.07 | 0.11 | 0.15 | 0.07 | 0.16 | 0.11 | 0.05 | 0.11 | 0.11 | 0.53 |
| Perch 2 | 0.03 | 0.05 | 0.09 | 0.00 | 0.01 | 0.05 | 0.01 | 0.04 | 0.05 | 0.36 |
| BirdCODE | **0.18** | **0.24** | **0.29** | **0.11** | **0.42** | **0.19** | **0.16** | **0.22** | **0.22** | **0.77** |

**Table S12.** Per-dataset Macro Recall (event@0.2): Results for each of the 68 WABAD datasets, average across WABAD, and results for XCSL. The best value in each column is shown in bold.

| Model | ARD | BAM | BIAL | BMT | BOLIN | BRCAS | BRE | BUR | CARI | CAT | CB | CLH | COU | CRUZ | DEVA |
|---|---|---|---|---|---|---|---|---|---|---|---|---|---|---|---|
| sl-BEATs-all | 0.05 | 0.03 | 0.24 | 0.04 | 0.04 | 0.04 | 0.05 | 0.03 | 0.04 | 0.22 | 0.11 | 0.32 | 0.20 | 0.03 | 0.06 |
| APN-20 | 0.10 | 0.04 | 0.29 | 0.06 | 0.08 | 0.08 | 0.10 | 0.06 | 0.10 | 0.25 | 0.12 | 0.36 | 0.22 | 0.10 | 0.09 |
| Perch 2 | 0.01 | 0.03 | 0.23 | 0.02 | 0.04 | 0.02 | 0.01 | 0.00 | 0.02 | 0.18 | 0.05 | 0.26 | 0.18 | 0.01 | 0.02 |
| BirdCODE | **0.23** | **0.08** | **0.40** | **0.14** | **0.17** | **0.27** | **0.23** | **0.16** | **0.23** | **0.45** | **0.22** | **0.54** | **0.29** | **0.21** | **0.24** |

| Model | DONG | DUNAS | DYOM | EMP | EVROS | FEU | FNCA | GLEN | GTLU | HAG | HAK | HAR | HONDO | HUAP | JUNCA |
|---|---|---|---|---|---|---|---|---|---|---|---|---|---|---|---|
| sl-BEATs-all | 0.02 | 0.16 | 0.05 | 0.23 | 0.21 | 0.06 | 0.03 | 0.25 | 0.01 | 0.09 | 0.00 | 0.13 | 0.15 | 0.04 | 0.19 |
| APN-20 | 0.03 | 0.23 | 0.10 | 0.37 | **0.26** | 0.04 | 0.08 | 0.35 | 0.05 | 0.14 | 0.01 | 0.18 | 0.18 | 0.16 | 0.23 |
| Perch 2 | 0.00 | 0.10 | 0.03 | 0.13 | 0.14 | 0.04 | 0.01 | 0.25 | 0.00 | 0.10 | 0.00 | 0.11 | 0.08 | 0.02 | 0.08 |
| BirdCODE | **0.09** | **0.47** | **0.15** | **0.39** | 0.24 | **0.19** | **0.13** | **0.50** | **0.12** | **0.21** | **0.14** | **0.22** | **0.23** | **0.29** | **0.29** |

| Model | KAR | KIB | LIM | MABI | MAPIMI | MARTI | MILLAN | MONTEB | MOPU | NAV | NL | OESF | OIO | OLIV | PETI |
|---|---|---|---|---|---|---|---|---|---|---|---|---|---|---|---|
| sl-BEATs-all | 0.26 | 0.02 | 0.00 | 0.32 | 0.17 | 0.07 | 0.06 | 0.06 | 0.09 | 0.18 | 0.10 | 0.40 | 0.03 | 0.27 | 0.13 |
| APN-20 | 0.33 | 0.07 | 0.08 | 0.21 | 0.20 | 0.10 | 0.18 | 0.18 | 0.14 | 0.17 | 0.25 | 0.40 | 0.12 | 0.19 | **0.27** |
| Perch 2 | 0.17 | 0.00 | 0.01 | 0.22 | 0.03 | 0.08 | 0.04 | 0.12 | 0.09 | 0.17 | 0.16 | 0.24 | 0.01 | 0.22 | 0.05 |
| BirdCODE | **0.50** | **0.14** | **0.12** | **0.37** | **0.40** | **0.20** | **0.23** | **0.19** | **0.24** | **0.32** | **0.32** | **0.55** | **0.31** | **0.42** | 0.24 |

| Model | PGF | PINA | PITI | POZO | PUUL | QR | RBA | RFP | RGU | RME | ROTOK | SAL | SBN | SCHF | SCHG |
|---|---|---|---|---|---|---|---|---|---|---|---|---|---|---|---|
| sl-BEATs-all | 0.01 | 0.14 | 0.01 | 0.04 | 0.03 | 0.19 | 0.12 | 0.16 | 0.26 | 0.19 | 0.03 | 0.08 | 0.09 | 0.49 | 0.11 |
| APN-20 | 0.00 | 0.24 | **0.23** | 0.03 | 0.06 | 0.19 | 0.17 | 0.30 | 0.38 | 0.39 | 0.01 | 0.12 | 0.06 | 0.38 | 0.14 |
| Perch 2 | 0.00 | 0.10 | 0.11 | 0.02 | 0.03 | 0.08 | 0.07 | 0.10 | 0.09 | 0.11 | 0.01 | 0.01 | 0.02 | 0.38 | 0.09 |
| BirdCODE | **0.05** | **0.38** | 0.22 | **0.18** | **0.14** | **0.41** | **0.37** | **0.50** | **0.51** | **0.49** | **0.13** | **0.21** | **0.24** | **0.70** | **0.16** |

| Model | SD | SITH | SLOB | SPMCO | TAM | UNI | VER | VIL | Avg | XCSL |
|---|---|---|---|---|---|---|---|---|---|---|
| sl-BEATs-all | 0.05 | 0.16 | 0.16 | 0.06 | 0.14 | 0.11 | 0.10 | 0.16 | 0.12 | 0.64 |
| APN-20 | 0.09 | 0.13 | 0.16 | 0.12 | 0.20 | 0.16 | 0.13 | 0.23 | 0.17 | 0.72 |
| Perch 2 | 0.02 | 0.10 | 0.14 | 0.01 | 0.04 | 0.06 | 0.03 | 0.08 | 0.08 | 0.50 |
| BirdCODE | **0.17** | **0.37** | **0.30** | **0.17** | **0.43** | **0.25** | **0.26** | **0.29** | **0.28** | **0.84** |

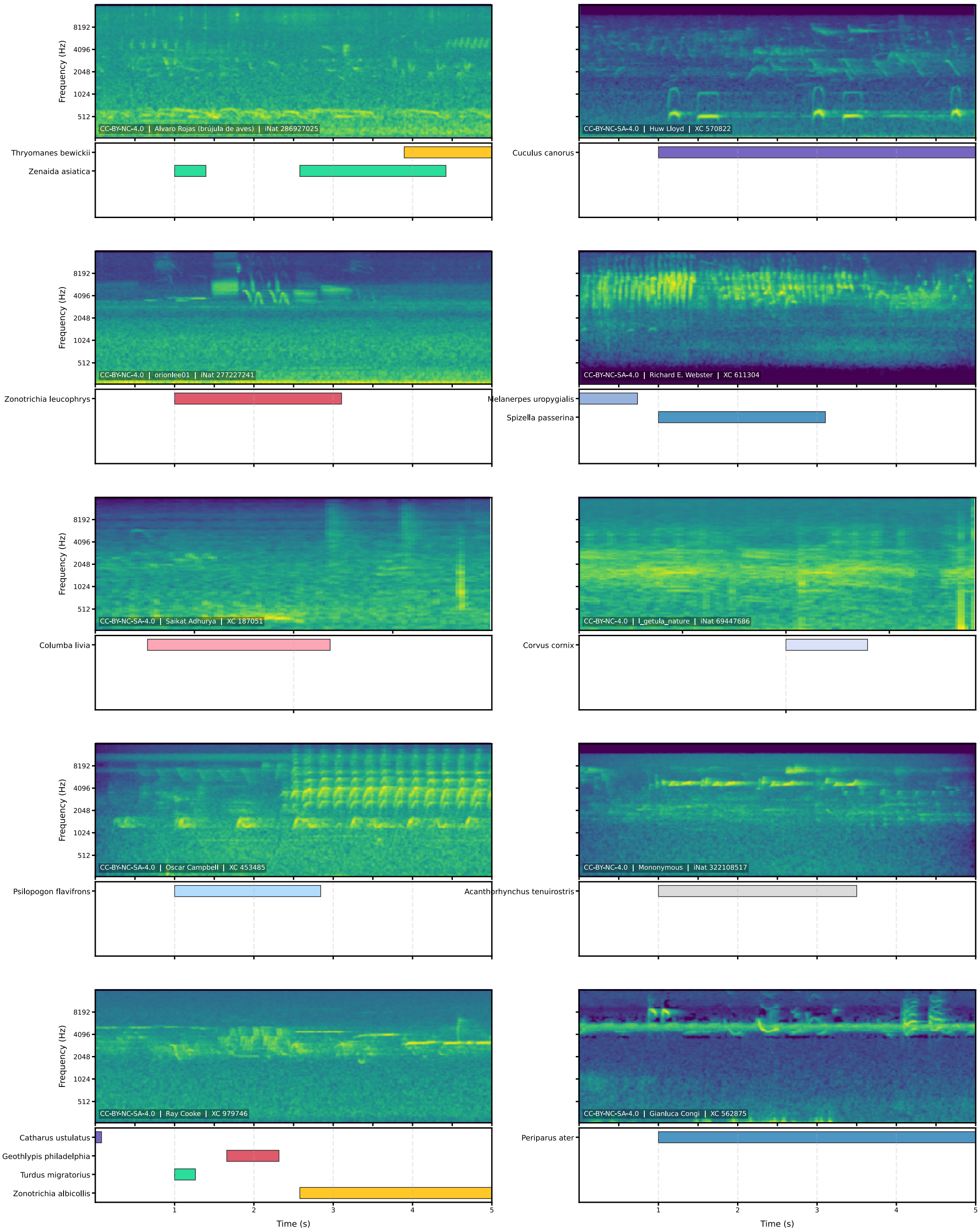


**Figure S2.** Example detections on xeno-canto and iNaturalist datasets.

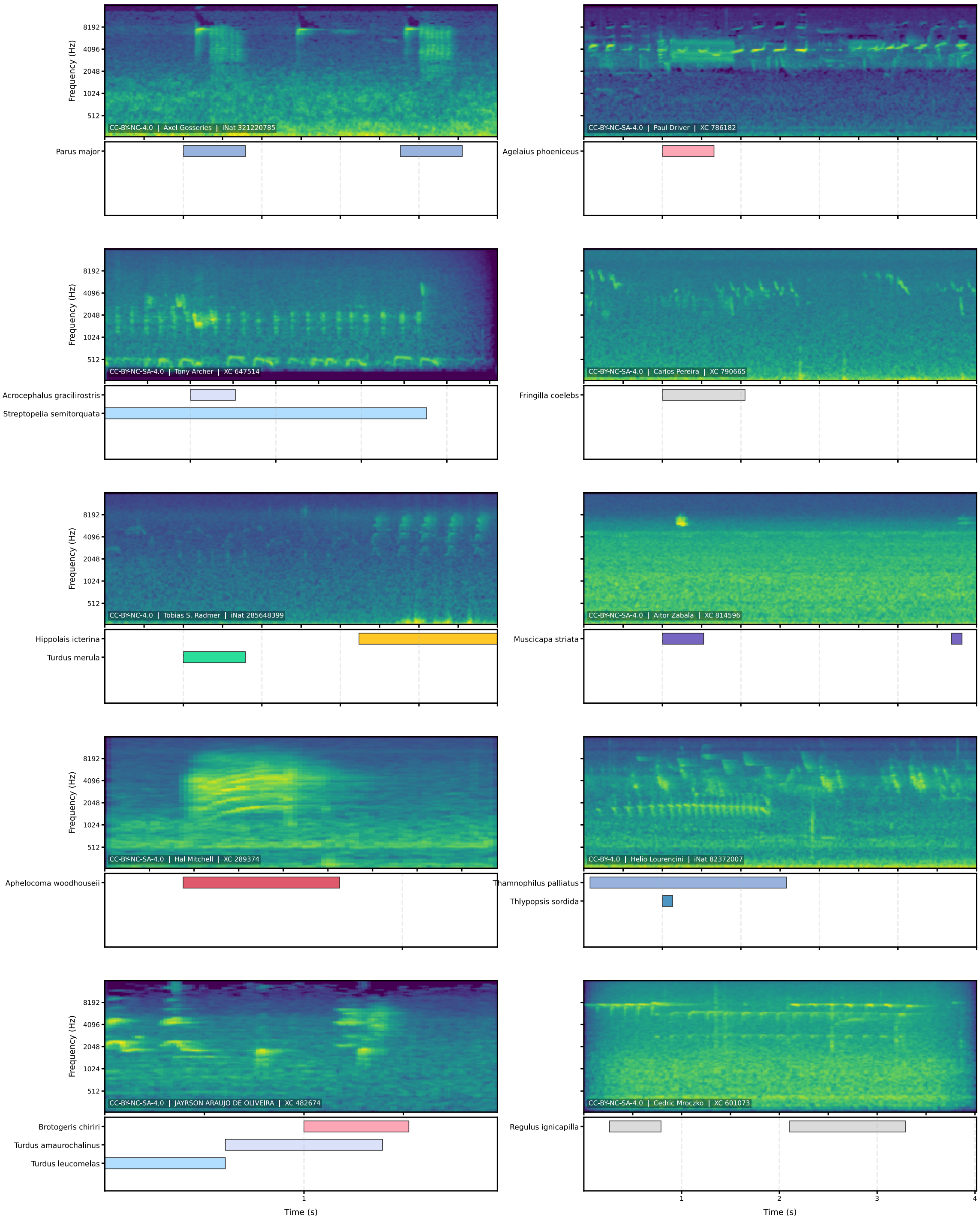


**Figure S3.** Example detections on xeno-canto and iNaturalist datasets.

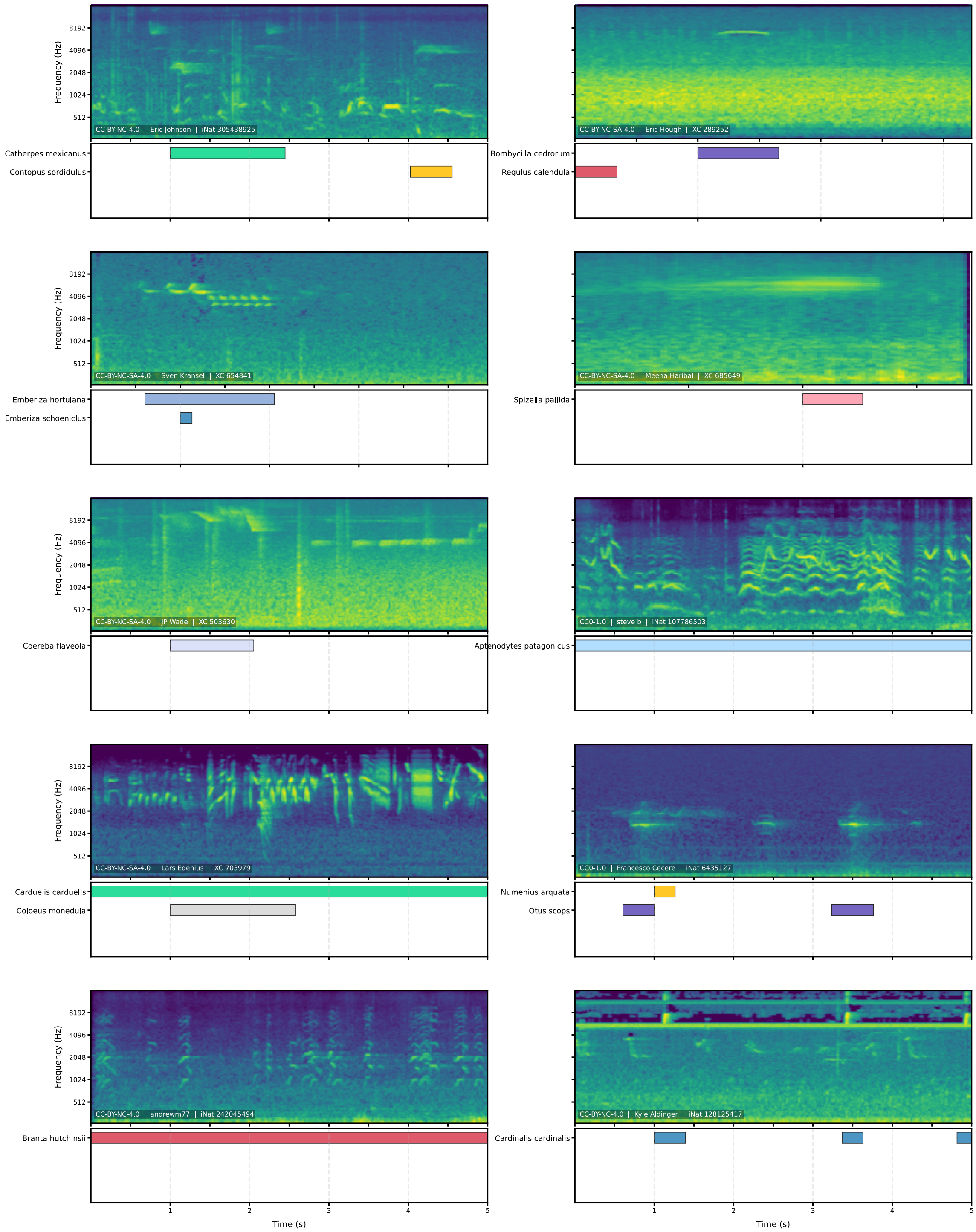


**Figure S4.** Example detections on xeno-canto and iNaturalist datasets.

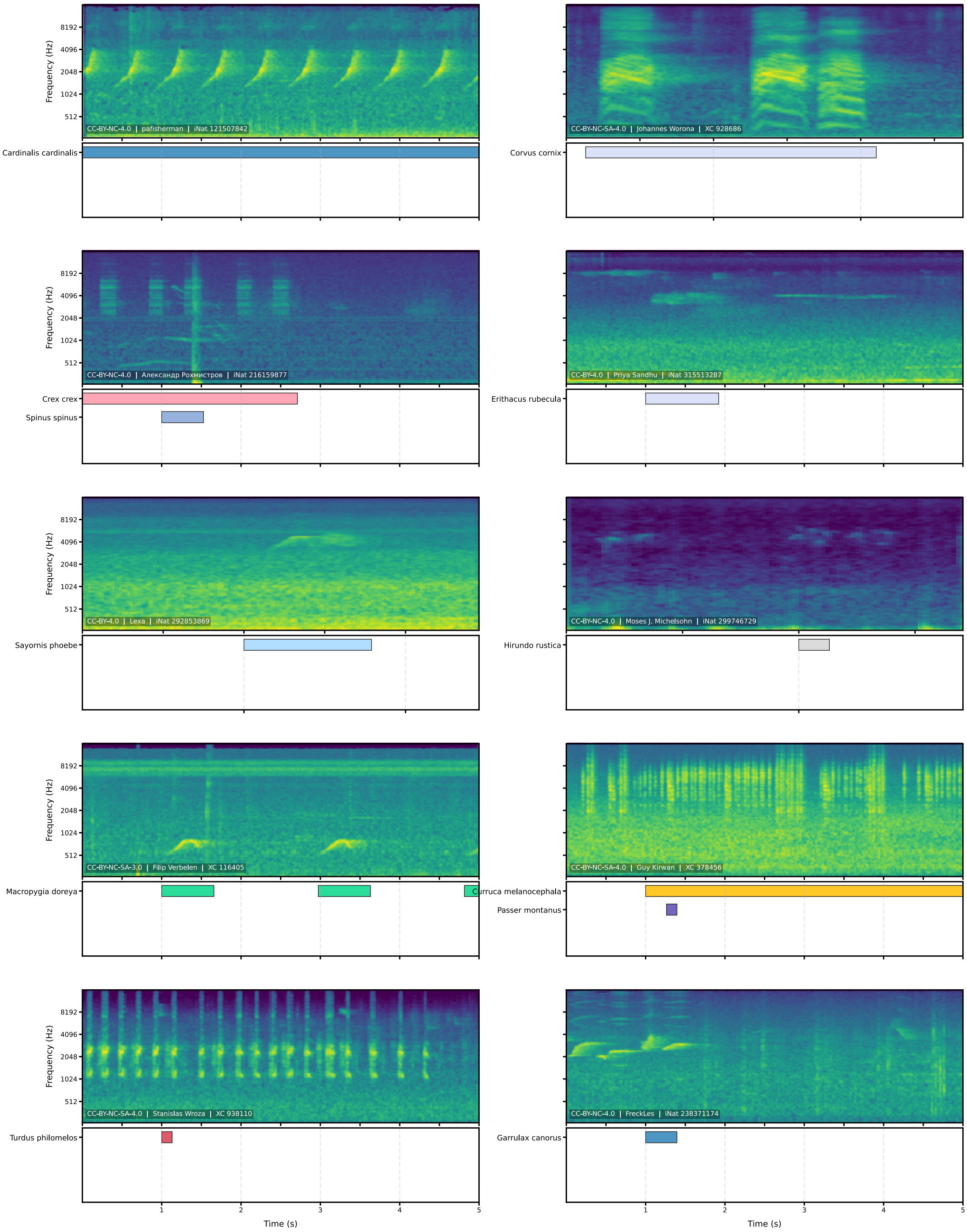


**Figure S5.** Example detections on xeno-canto and iNaturalist datasets.

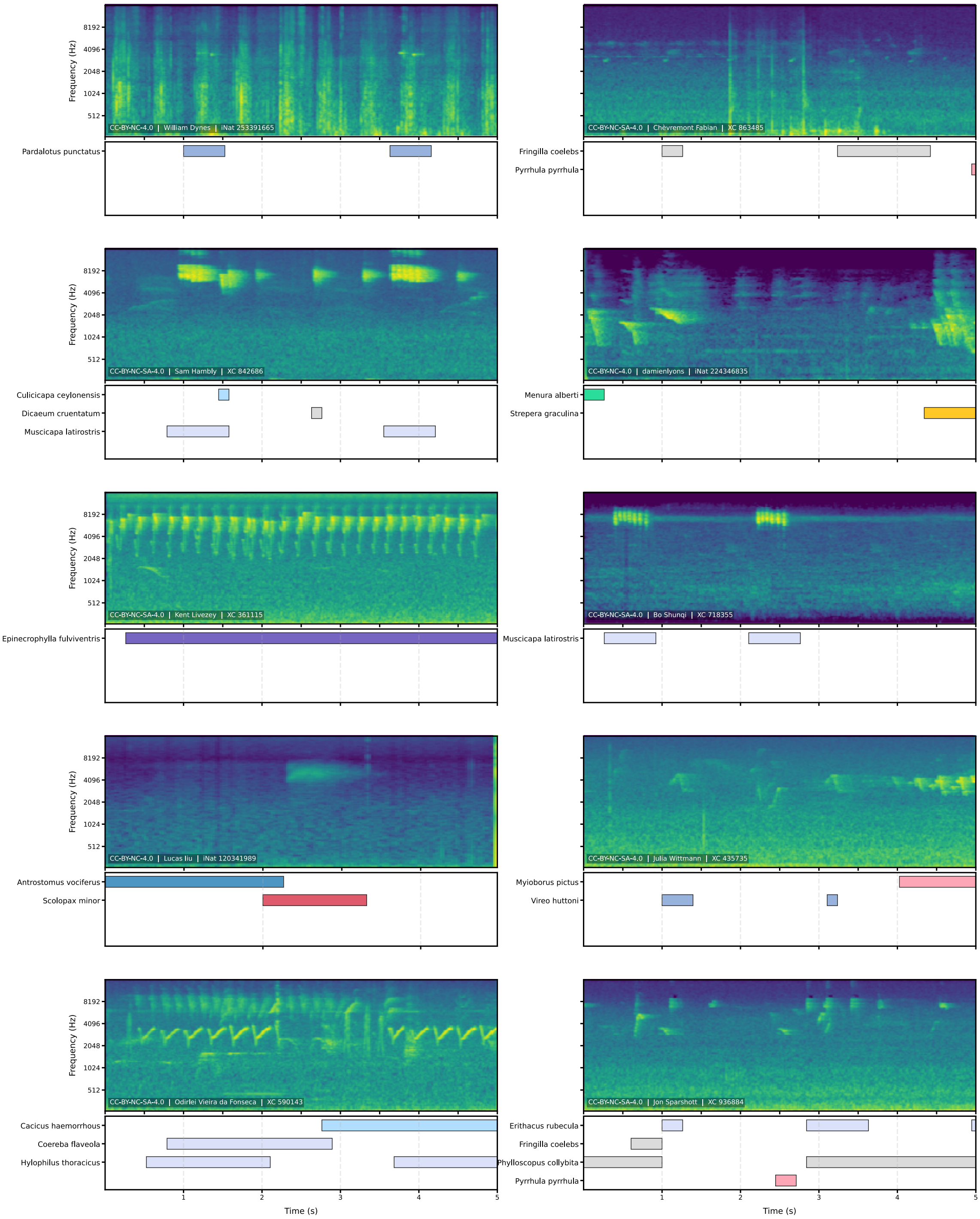


**Figure S6.** Example detections on xeno-canto and iNaturalist datasets.
